# Dynamic modulation of spleen germinal center reactions by gut bacteria during *Plasmodium* infection

**DOI:** 10.1101/2021.02.02.429404

**Authors:** Rabindra K. Mandal, Joshua E. Denny, Ruth Namazzi, Robert O. Opoka, Dibyadyuti Datta, Chandy C. John, Nathan W. Schmidt

## Abstract

Gut microbiota educate the local and distal immune system in early life to imprint long-term immunological outcomes while maintaining the capacity to dynamically modulate the local mucosal immune system throughout life. It is unknown if gut microbiota provide signals that dynamically regulate distal immune responses following an extra-gastrointestinal infection. Using the murine model of malaria, we show that existing spleen germinal center reactions are malleable to dynamic cues provided by gut bacteria that impact parasite burden. Gut bacteria composition was also shown to correlate with the severity of malaria in humans. Whereas antibiotic-induced changes in gut bacteria has been associated with immunopathology or impairment of immunity, our data demonstrate antibiotic-induced changes in gut bacteria can enhance humoral immunity to *Plasmodium*. This effect is not universal, but depends on baseline gut bacteria composition. These data demonstrate the dynamic communications that exist between gut bacteria and the systemic immune system as well as the plasticity of an ongoing humoral immune response.

**Summary:** The study by Mandal R, et al. provides new insight into the dynamic communications that exist between gut bacteria, the systemic immune system and the plasticity of spleen germinal center reactions during *Plasmodium* infection.

## Introduction

Gut microbiota, the community of microorganisms living in the gastrointestinal tract, have diverse effects on host biology. One important role of gut microbiota in early life is to educate the mucosal and systemic immune system to reduce the risk of immunopathology that may occur later in life. In mice, neonatal exposure to gut bacteria protects adult mice against experimental inflammatory bowel disease (IBD) via exposure to sphingolipids that inhibit the accumulation of mucosal iNKT (An et al., 2014; Olszak et al., 2012). Early life exposure to high-diversity gut microbiota also prevents elevated serum IgE in mice and reduces the risk of oral-induced anaphylaxis (Cahenzli et al., 2013). The reported increase in serum IgE after weaning is consistent with gut microbiota composition in mice programming the immune system during weaning through induction of regulatory T cells to resist inflammatory pathologies into adulthood (Al Nabhani et al., 2019). Perturbations of the microbiota during this critical “weaning window” alter immune education that result in increased susceptibility to immunopathologies (colitis, allergic inflammation, and cancer) into adulthood (Al Nabhani et al., 2019), and reduced responsiveness in adolescent mice to numerous adjuvanted and non-adjuvanted vaccines (Lynn et al., 2018). Gut microbiota-dependent education of the immune system also occurs in humans. The type of LPS present in early life, which is influenced by gut bacteria composition, can educate the immune system to be resistant or susceptible to autoimmune diseases, including type 1 diabetes, later in life (Vatanen et al., 2016). Additionally, events in early life that can alter gut microbiota composition (*e.g*., caesarean section, formula feeding, infection/inflammation, and antibiotic exposure) can pathologically imprint the immune system and increase the risk of childhood asthma and atopy (Arrieta et al., 2015; Fujimura et al., 2016; Shao et al., 2019). Consistent with gut microbiota educating or priming the host immune system to respond in a prescribed nature, mice treated with antibiotics 2-4 weeks prior to viral infection (influenza or LCMV) or administration of the seasonal influenza vaccine had reduced adaptive immune responses (Abt et al., 2012; Ichinohe et al., 2011; Oh et al., 2014) resulting in impaired control of the virus (Abt et al., 2012; Ichinohe et al., 2011). It has also been shown that humans treated with antibiotics prior to administration of the seasonal influenza vaccine exhibited reduced antibody titers to one of three viral strains present in the vaccine (Hagan et al., 2019). Although antibiotics are extremely beneficial at eliminating pathogenic bacteria, these data support the idea that antibiotic-induced changes to gut bacteria populations can be detrimental to host immunity.

The interaction between gut microbiota and intestinal mucosal immune system is also dynamic and extends beyond early life (Wang and Li, 2019). For example, imbalanced gut microbiota composition leads to altered interactions between these gut microbiota and the host gut mucosal immune system, and as a consequence the host becomes susceptible to bacterial translocation, gut-derived infection, and detrimental clinical outcomes like IBD, autoimmunity and allergy (Khan et al., 2019; Round and Mazmanian, 2009; Schuijt et al., 2013). However, it is not known if gut microbiota also provide dynamic modulation of a systemic immune response to an extra-intestinal infection.

*Plasmodium* species are the causative agent of malaria and were responsible for 228 million episodes of malaria and 405,000 deaths in 2018 (WHO, 2019). *Plasmodium* infections are initiated when an infected mosquito delivers sporozoites during a blood meal. Sporozoites infect hepatocytes, initiating the liver stage infection, where they differentiate into merozoites that are released into circulation and establish cyclical infection of red blood cells (Cowman et al., 2016). The blood stage infection is controlled by the formation of germinal center (GC) reactions that result in the production of high-affinity antibodies that mediate clearance of the parasite (Figueiredo et al., 2017; Guthmiller et al., 2017; Obeng-Adjei et al., 2015; Pérez-Mazliah et al., 2015; Pérez-Mazliah et al., 2017; Ryg-Cornejo et al., 2016).

Recent research has identified a connection between gut microbiota and malaria. For example, gut microbiota can induce cross-reactive antibodies that confer protection against sporozoites, although these antibodies did not provide protection against *Plasmodium* blood stage infection (Yilmaz et al., 2014). Another study, in which Malian children were longitudinally tracked from the end of the dry season (i.e. no malaria transmission) through the ensuing malaria transmission season, demonstrated a correlation between stool bacteria composition at the end of the dry season and the rate at which children were infected with *Plasmodium falciparum* (Yooseph et al., 2015), for ethical reasons this report did not assess severe malaria. Finally, the composition of gut microbiota has been shown to shape the severity of malaria in mice (Denny et al., 2019; Stough et al., 2016; Villarino et al., 2016). Numerous knowledge gaps remain in understanding malaria-gut microbiome interactions. It is unknown which of the gut microbiota constituents (bacteria, viruses, fungi, etc.) affect the severity of malaria. It is not known if the gut microbiome affects the severity of malaria in humans. Finally, it is unclear how gut microbiota interacts with the host immune system to ultimately impact parasite burden and the severity of malaria.

Here, we demonstrate that, among gut microbiota constituents, gut bacteria modulate the severity of malaria in mice. Importantly, we also report substantial differences in stool bacteria composition between Ugandan children with asymptomatic *Plasmodium falciparum* infection compared to severe malarial anemia. We show in mice that gut bacteria provide cues that dynamically modulate spleen GC reactions, which ultimately impact parasite burden, providing a potential explanation for the protection from severe malaria associated with stool bacteria compositions in the Ugandan children. Collectively, these data provide new insight into the dynamic communications that exists between gut bacteria and the systemic immune system and show that the effect of antibiotic-induced changes in gut bacteria populations on host immunity depends on the baseline bacteria composition. These data also support the possibility that human gut bacteria may impact the severity of malaria in children.

## Results

### Gut bacteria are mediators of gut microbiota-dependent modulation of malaria

To gain insight into how gut microbiota modulate the severity of malaria and host immunity, reciprocal ceca content transplants (Fig. S1A) were performed between C57BL/6 mice that are either resistant (Taconic Biosciences; Tac) or susceptible (Charles River Laboratories; CR) to developing hyperparasitemia and severe anemia following *Plasmodium yoelii* 17XNL infection (Villarino et al., 2016). Control Tac and CR mice that received ceca contents from Tac and CR mice, respectively, exhibited the expected low (Tac→Tac) and high (CR→CR) parasite burden following *P. yoelii* 17XNL infection (Fig. S1B-C). CR ceca content transplants into Tac mice (CR→Tac) resulted in the Tac mice developing a high parasite burden similar to CR control mice, whereas Tac ceca content transplants into CR mice (Tac→CR) did not reduce their elevated parasite burden (Fig. S1B-C). 16S rRNA gene sequence analysis was done by Multiple 16S Variable Region Species-Level IdentificatiON (MVRSION) to analyze gut bacteria populations (Schriefer et al., 2018). Consistent with the parasite burden data, the Tac→Tac mice displayed different bacteria communities than the CR→Tac, Tac→CR and CR→CR mice (Fig. S1D-E). Of note, the Tac and CR donor bacteria communities differentiate from the Tac→Tac or CR→CR samples because they were from the ceca contents versus fecal pellets, respectively. These data demonstrated that the bacteria in CR mice are ecologically dominant over the bacteria in Tac mice. They also revealed that while ceca content transplants from CR→Tac mice provide an approach to probe how gut microbiota affect host immunity in Tac mice, they do not allow for manipulation of gut microbiota-host immunity interactions in CR mice.

Currently, it is not known which constituent of gut microbiota (fungi, virus, bacteria, etc.) affects the severity of malaria. To dissect the contribution of these constituents on *P. yoelii* parasite burden, Tac and CR mice were treated with antifungals, antivirals, or antibiotics in drinking water to modulate those gut microbes. Antifungal treatment in CR mice had minimal effects on *P. yoelii* parasite burden, whereas Tac mice treated with 5-fluorocytosine showed an increase in parasite burden (Fig. S2A-D), suggesting that 5-fluorocytosine-sensitive fungi may contribute to the hyperparasitemia resistant phenotype. Yet, toxicity associated with these treatments, in particular the antifungal cocktail, and inconsistent effects of 5-fluorocytosine treatment on parasitemia in Tac mice over multiple experiments made it difficult to interpret those results. Antiviral treatment resulted in a slight decrease in the peak parasite burden in CR mice, while no effect in parasite burden in Tac mice (Fig. S2E-F). Collectively, these data suggested that gut fungi and viruses targeted by these antimicrobials minimally modulate the severity of malaria following *P. yoelii* infection.

To assess the contribution of gut bacteria towards severity of malaria, mice were treated with one of five different antibiotics in the drinking water for 14-days before and continuously after infection with *P. yoelii* (Fig. 1A-B). Each of the antibiotics resulted in a decrease in parasite burden in CR mice compared to the control CR mice, with ampicillin, gentamicin, metronidazole, and vancomycin treatment exerting the greatest effect (Fig. 1C-D). In the Tac mice, the effect of antibiotic treatment on *P. yoelii* parasite burden was less pronounced, but those antibiotic treatments that did have an effect (ampicillin, gentamicin, and vancomycin) trended towards increased parasite burden compared to the control Tac mice (Fig. 1E-F), which was in contrast to the decreased parasite burden that was observed in antibiotic treated CR mice. Given that gentamicin and vancomycin have little to no intestinal absorption (Fig. 1B) (Koga et al., 2006; Rao et al., 2011), and that treatment with ampicillin, gentamicin, and vancomycin resulted in parasite burden shifting in opposite directions compared to the control CR and Tac mice, respectively, these data suggest that these antibiotics did not have a direct effect on *P. yoelii*, rather they exerted an indirect effect through changes in gut bacteria composition.

**Figure 1.**
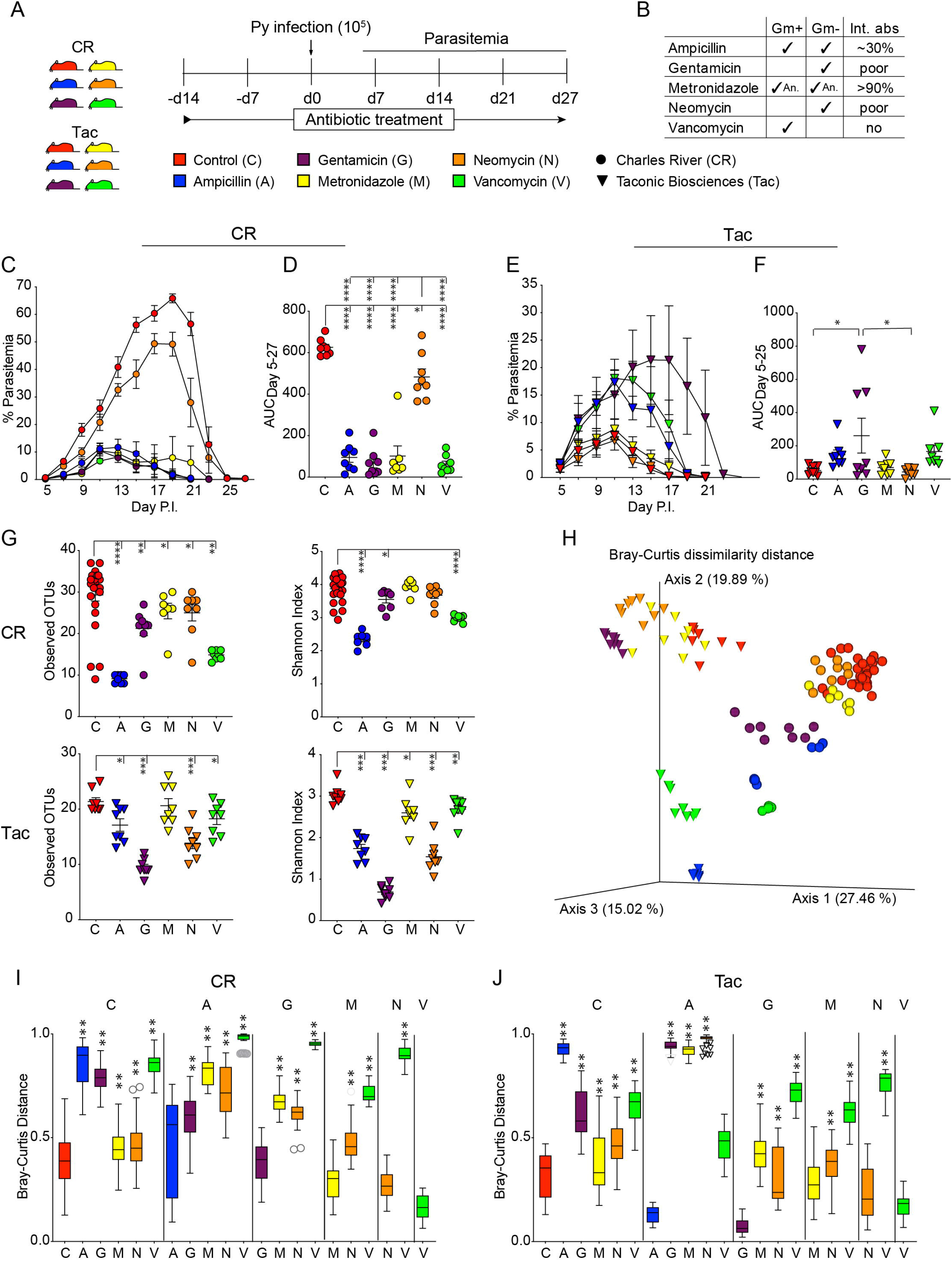
Bacteria are a critical gut microbiota constituent that modulate the severity of malaria. (**A**) C57BL/6 mice from Charles River (CR) and Taconic (Tac) mice were treated with one of the five antibiotics in drinking water from two weeks prior to infection till the clearance of infection. Mice were infected with *Plasmodium yoelii* 17XNL on day 0 and parasitemia was tracked on every other day from day 5 post infection (p.i.) as indicated. (**B**) Spectrum of antibiotics against bacteria and intestinal absorption (Int. abs). Gm+: Gram positive bacteria; Gm-: Gram negative bacteria; An: Anaerobic. (**C** and **D**) Parasitemia and AUC of CR mice. (**E** and **F**) Parasitemia and AUC of Tac mice (**G-J**) Gut microbiota analysis from fecal pellets at day 0 using MVRSION (Multiple 16S Variable Region Species-Level IdentificatiON). (**G**) Alpha diversity analysis of CR and Tac. (**H**) Beta diversity analysis of CR and Tac using Bray-Curtis distance. (I and J) Bray-Curtis distance comparison between groups of CR and Tac mice respectively. Box end shows lower and upper quartile and horizontal line inside box is median. Y-axis shows Bray-Curtis dissimilarity distance of groups on X-axis to groups on the top of vertical columns. Data are means ± SEM. Statistical significance were analyzed by (**D** and **F**) one-way ANOVA with Tukey’s multiple comparisons test, (**G**) pairwise Kruskal-Wallis test, and (**I** and **J**) pairwise PERMANOVA with 999 permutations. Significance level is compared between group at the top of column to group on X-axis. (**G**) Only comparisons with control are shown. Data are cumulative from 2 independent experiments with n=4 in each trial. * = *p* < 0.05, ** = *p* < 0.01, **** = *p* < 0.0001.

MVRSION analysis of gut bacteria populations identified that each antibiotic treatment changed the richness and abundance of bacteria species (alpha diversity) as measured by both observed operational taxonomic units (OTUs; defined at 97% sequence identity) that measures species richness and Shannon index that measures both species richness and their abundance (Fig. 1G), and composition of the bacteria populations (beta diversity) as measured by the Bray-Curtis dissimilarity distance (Fig. 1H) compared to the control CR and Tac mice. Bacteria populations were also significantly different between each of the individual antibiotic treated groups within CR mice (Fig. 1I) and Tac mice (Fig. 1J). A supervised machine learning model was able to classify *Plasmodium* susceptibility phenotype (high versus low parasitemia) in mice with 97% overall accuracy (Fig. S3A-C). Metronidazole-treated mice were excluded from the model owing to the inability to attribute decreased parasitemia to antibiotic-induced changes in gut microbiota versus the known *anti-Plasmodium* effects of metronidazole (James, 1985).

Intriguingly, differential abundance of just 8 bacterial species were able to classify malaria susceptibility in these mice (Fig. S3D-E). Collectively, these data suggest that gut bacteria can profoundly impact the severity of malaria in mice. Moreover, they identify antibiotic treatments as an effective tool to probe how gut bacteria affect severity of malaria in CR mice. In particular, vancomycin treatment is useful in this model system given the pronounced effect of treatment on bacteria populations and the inability of vancomycin to be absorbed through the intestinal tract, thereby limiting the observed effect of vancomycin treatment on *P. yoelii* parasite burden to the effect of this antibiotic on gut bacteria. These data also suggest that the effect of antibiotic-induced changes in gut bacteria compositions on host immunity is not always detrimental, and in fact this effect can actually be beneficial for host protection. The outcome is likely dependent on the baseline gut bacteria composition and nature of the infection or vaccine.

### Stool bacteria populations correlate with severity of malaria in Ugandan children

To determine if gut bacteria composition is associated with the severity of malaria in humans, bacteria populations were analyzed by MVRSION in stool collected from Ugandan children who had severe malaria anemia (SMA; n=40), or community control children who were otherwise healthy, but had asymptomatic *Plasmodium falciparum* (Pf) infections (Pf Pos; n=7) or were Pf negative (Pf Neg; n=28) (Fig. 2A). Although there was a relatively low number of samples of children with asymptomatic Pf parasitemia (n=7), the bacteria species accumulation curve showed a plateauing in the curve within this group (Fig. S4), suggesting that additional samples may not have dramatically increased the total number of bacterial species identified in this group. There was no difference in the observed OTUs, Shannon index and pielou_e between children based on categorical variables of interest as antibiotic usage, enrollment site and sex except *Plasmodium* status (Fig. S5A). There were no significant linear correlations (−0.9 > r > 0.9 and *p* < 0.05) between the various continuous covariates of interest that were investigated (Fig. S5B). There was also no significant association in observed OTUs, Shannon index or pielou_e and any of the continuous covariates except for age (Fig. S5C). Pielou_e was negatively correlated with age (Fig. S5D). In contrast to prior publication (Bokulich et al., 2016) but similar to our previous publication in Kenyan infants (Mandal et al., 2018), there was no increase in alpha diversity, measured by observed OTUs or Shannon index, as age increased (Fig. S5C). There was no difference in observed OTUs based on *Plasmodium* status of children (Fig. 2B). Shannon index was significantly different between Pf Neg vs Pf Pos samples, while Pielou_e was significantly different between Pf Neg vs Pf Pos and Pf Pos vs SMA (Fig. 2C). These data show that there were a few differences in stool bacteria alpha diversity based on *Plasmodium* status of children.

**Figure 2.**
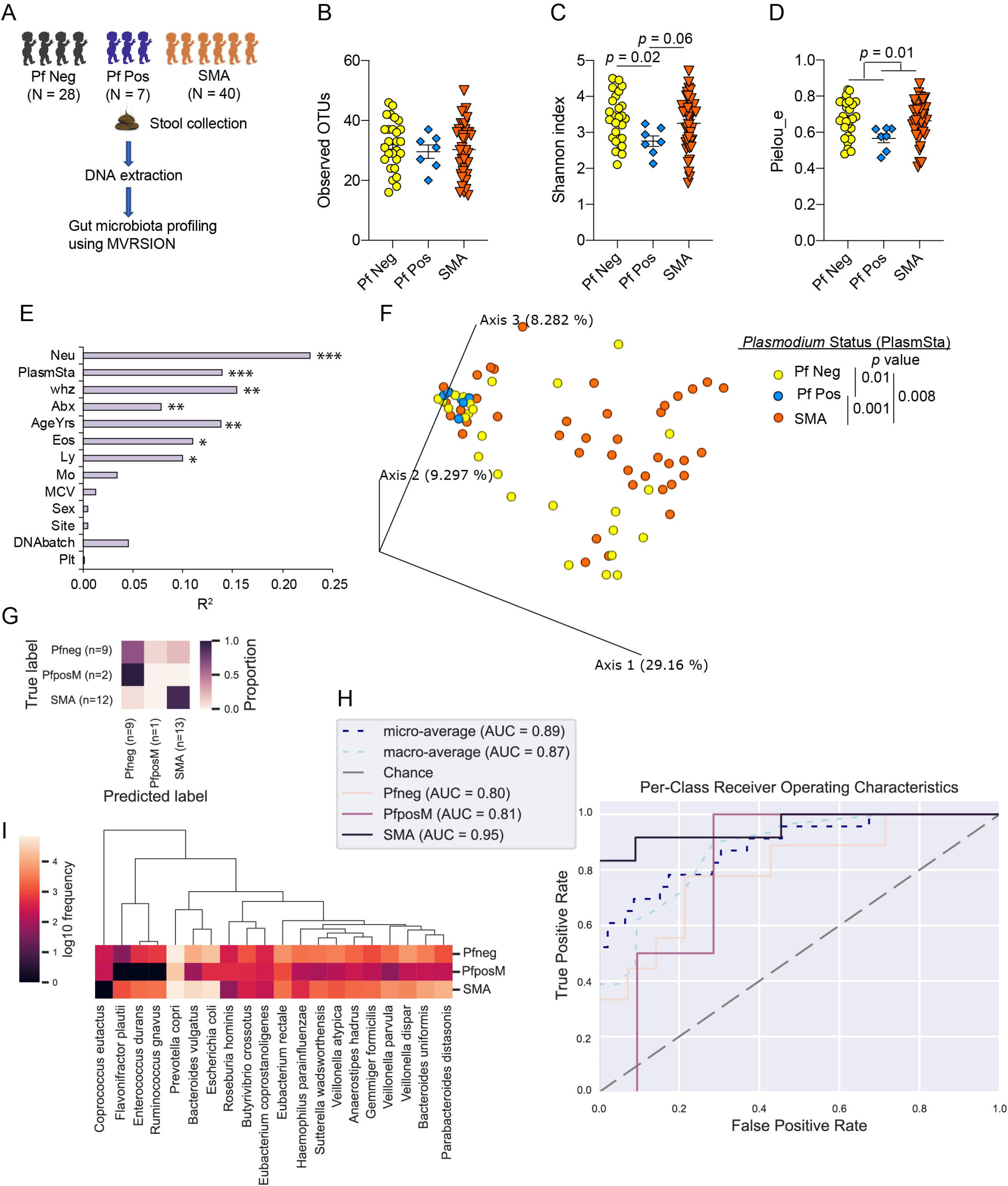
Gut microbiota composition is associated with severity of malaria in humans. (**A**) Stool samples were collected from children at the time of hospitalization diagnosed with severe malaria anemia (SMA, n=40) prior to antimalarial treatment and community control children, Pf negative (Pf Neg, n=28) and asymptomatic Pf positive (Pf Pos, n=7) at the time of enrollment and stored at −80°C. Stools were subjected to gut microbiome analysis using 16S rRNA gene sequencing using MVRSION. One Pf Neg sample was removed from alpha and beta diversity analysis due to low sequencing depth. (**B-D**) Alpha diversity analysis of *Plasmodium* status measured using Observed OTUs, Shannon index and Pielou_e respectively. (**E**) Beta diversity variation explained by different co-variates using Bray-Curtis distance matrix. (**F**) Principal coordinate analysis (PCoA) plot shows clustering of based on *Plasmodium* status using Bray-Curtis distance. (**G-I**) Predicting categorical sample (PlasmSta) with supervised machine-learning classifiers using random forest. (**G**) Result of prediction with test set data. (**H**) Per-class receiver operating characteristics curve. (**I**) Top 20 species that are the most predictive of the three groups. Data are means ± SEM. The following statistical tests were used to analyze the data; pairwise Kruskal-Wallis test (**B-D**), “envfit” function of vegan package implemented in R (**E**), and “adonis” function of vegan package implemented in R accounting for the covariates that significantly explained the variation of gut microbiota composition in Fig. 2E (**F**). PlasmSta – Plasmodium status (SMA, Pf Pos, Pf Neg); whz - weight for height z-score; Neu, Eos, Ly, Mo, Plt – Absolute count (K/mcl) of blood parameters neutrophils, eosinophils, lymphocytes, monocytes, and platelets, respectively; Abx – Antibiotic use; AgeYrs – Age in years; Site – Study area; Sex – Sex of participants; DNAbatch – Batch of DNA extraction; MCV – mean corpuscular volume (fL). * = *p* < 0.05, ** = *p* < 0.01.

Stool bacterial beta diversity (bacterial composition compared between samples) was also evaluated between the Pf Neg, Pf Pos and SMA groups. The “envfit” function (vegan package) identified seven covariates that explained significant variation in beta diversity measured by the Bray-Curtis dissimilarity distance. These included, absolute neutrophil count, *Plasmodium* status, weight for height Z-score (whz), antibiotic usage, age, absolute eosinophils and lymphocyte count (Fig. 2E and Fig. S6). Stool bacteria composition was significantly different between Pf Neg vs Pf Pos (*p* = 0.01), Pf Neg vs SMA (*p* = 0.008) and Pf Pos vs SMA (*p* = 0.001, ANOVA) measured by Bray-Curtis dissimilarity distance accounting for all the seven covariates (Fig. 2F). Age and antibiotics are well known to affect the gut microbiota composition (Hasan and Yang, 2019). However, there was no significant interaction among *Plasmodium* status, age of child and antibiotic usage (*p >* 0.05, ANOVA; Fig. S7). Of note, Pf Pos versus SMA were more significantly different compared to Pf Neg vs SMA and Pf Neg vs Pf Pos (Fig. 2F). Although, stool samples from SMA children were collected at varying days (0-22) after hospitalization and enrollment, gut bacteria populations measured by beta and alpha diversity were not different between the varying days (Supplementary Table 1; Fig. S7B-C). This provides support that the differential stool bacteria populations observed in SMA children compared to the Pf Pos children were not simply a product of the severe infection causing changes in gut bacteria. Further supporting an association between gut bacteria compositions and severity of Pf infections, a supervised machine learning model using random forest was able to predict the *Plasmodium* status with 74% overall accuracy (Fig. 2G-H and Fig. S8A). Both higher and lower relative abundant bacterial species were predictive of *Plasmodium* status (Fig. 2I and Fig. S8B). These data identify that there are different stool bacteria populations between these Ugandan children that have an asymptomatic *P. falciparum* infection compared to children that have SMA.

### Vancomycin treatment causes long-term changes in adult mouse gut bacteria populations that affect severity of malaria

Gut microbiota in infancy are capable of conferring long-term imprinting of the host immune system (Yang et al., 2016). It has also been shown that gut microbiota engage the mucosal immune system in a dynamic manner (Wang and Li, 2019). In the previous experiment (Fig. 1), mice were continuously treated with antibiotics, which precluded the ability to assess whether changes in gut bacteria before or after the *P. yoelii* infection resulted in the changes in severity of malaria. To determine the extent of gut microbiota imprinting on the host immune response to *Plasmodium* infection, groups of adult CR mice were provided vancomycin drinking water 4-weeks before *P. yoelii* infection and treated for at least 2-weeks. Vancomycin treatment was stopped 14 days before, 7 days before, 3 days before, or the day of *P. yoelii* infection (Fig. S9A). Regardless of when vancomycin treatment ceased, the vancomycin-treated CR mice displayed significantly lower parasitemia than the control CR mice (Fig. S9B-C). These data suggest that the early life gut microbiota do not imprint programmed immunological responses to *Plasmodium* that are unchangeable, but rather that the gut microbiota may imprint a particular immune response to *Plasmodium* infection when gut microbiota are modulated up to the day of *P. yoelii* infection. Moreover, the 14-day cessation in vancomycin before *P. yoelii* infection provided further support that the effect of vancomycin treatment on parasitemia is not attributed to a direct effect of vancomycin in *P. yoelii*, but rather vancomycin-induced changes in gut bacteria compositions.

The observation that a 14-day vancomycin treatment followed by 14 days of no treatment resulted in a significant decrease in parasitemia led to the question, how long does the protection afforded by antibiotic treatment last after cessation of treatment? To probe this question, CR mice were treated with vancomycin for 14 days and then placed on regular drinking water for either 90, 60, 30, or 15 days before *P. yoelii* infection (Fig. 3A). Strikingly, mice that were treated with vancomycin for 14 days and then rested for 90 days showed a profound decrease in parasitemia to levels that were barely detectable, which was in contrast to the age-matched control mice that developed the expected high levels of parasitemia (Fig. 3B-C).

**Figure 3.**
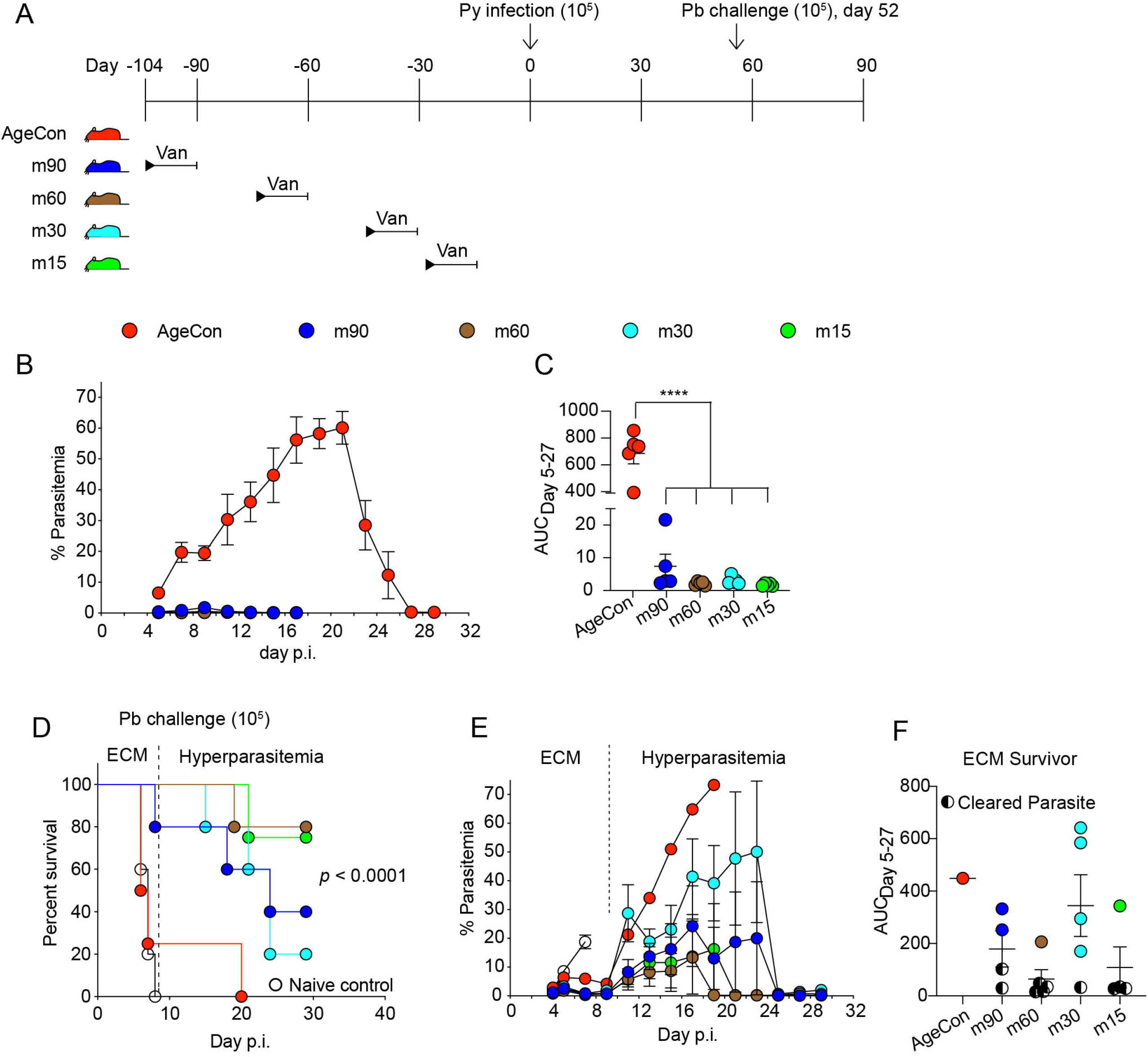
Vancomycin treatment provides protection against *P. yoelii* for months and imparts cross-species immunity to experimental cerebral malaria. (**A**) C57BL/6 mice from CR (susceptible) were treated with vancomycin for two weeks and put back on regular drinking water for 90 (m90), 60 (m60), 30 (m30), and 15 (m15) days before *P. yoelii* infection. Mice were challenged with *P. berghei* ANKA on day 52 post-infection. All of the mice were procured at the same time. (**B** and **C**) Parasitemia and AUC of *P. yoelii* infection. (**D**) Survival curve after *P. berghei* ANKA challenge. (**E** and **F**) Parasitemia and AUC after *P. berghei* ANKA challenge. Data are means ± SEM. (**C**) one-way ANOVA with Tukey’s multiple comparisons test. (**D**) Pairwise Log-rank (Mantel-Cox) test. Representative of two experiments with 5 mice per group in each trial. **** = *p* < 0.0001. Similar results were obtained from two independent experiments.

*P*. yoelii-immune mice confer cross-species protection against an otherwise lethal *Plasmodium berghei* ANKA infection that causes experimental cerebral malaria (ECM) or in the rare ECM-survivor, hyperparasitemia and severe anemia (Kurup et al., 2017). The extremely low *P. yoelii* parasite burden in the vancomycin treated mice led to the question of whether these mice elicited an adaptive immune response that would provide protection against *P. berghei* ANKA infection. To test this possibility, each of these groups of CR mice, along with age-matched *P. yoelii*-naïve CR mice, were challenged with *P. berghei* ANKA 52 days after the original *P. yoelii* infection (Fig. 3A). All of the naïve control mice died of ECM within 10 days of infection (Fig. 3D), and *P. yoelii*-naïve vancomycin treated CR mice were also susceptible to ECM (Fig. S9D). Whereas vancomycin control *P. yoelii-immune* CR mice remain susceptible to ECM, all of the groups of CR mice that had been treated with vancomycin and infected with *P. yoelii* exhibited high rates of survival from ECM (Fig. 3D). Moreover, among all of the *P. yoelii-*immune, ECM-survivor, vancomycin treated mice (minus 90, 60, 30, and 15 days), 56% (10 of 18) were also protected from *P. berghei* hyperparasitemia (Fig. 3E-F).

Antibiotic treatment in early life can have long-term effects on gut bacteria (Nobel et al., 2015). In adult populations, the long-term effect of antibiotic treatments on gut bacteria populations is less clear. Some reports have noted a recovery of baseline gut bacteria within 4 weeks post-treatment cessation (Palleja et al., 2018; Suez et al., 2018), whereas others have observed differences that last many months after antibiotic cessation (Haak et al., 2018). To determine if the long-term effect of 2-week vancomycin treatments on *P. yoelii*-parasite burden correlated with sustained changes in gut bacteria populations, bacteria community analysis was performed. MVRSION analysis of fecal pellets from age-matched control and vancomycin-treated CR mice (Fig. 3A) collected prior to *P. yoelii* infection on day 0 p.i. revealed a loss in alpha diversity in the vancomycin-treated mice that lasted up to 90 days after vancomycin treatment had stopped (Fig. 4A-B). Bacterial community composition, as measured by Bray-Curtis dissimilarity distance, demonstrated clear differences in bacteria populations between any of the vancomycin treated mice and the control CR mice (Fig. 4C-D), with all but two of the minus 90 day mice tightly clustered together with minimal differences between any of the vancomycin-treated mice compared among one another (Fig. 4D). Collectively the data indicate within the context of a *Plasmodium* infection that the composition of gut bacteria at the time of infection, rather than during early life, will ultimately impact the subsequent severity of infection.

**Figure 4.**
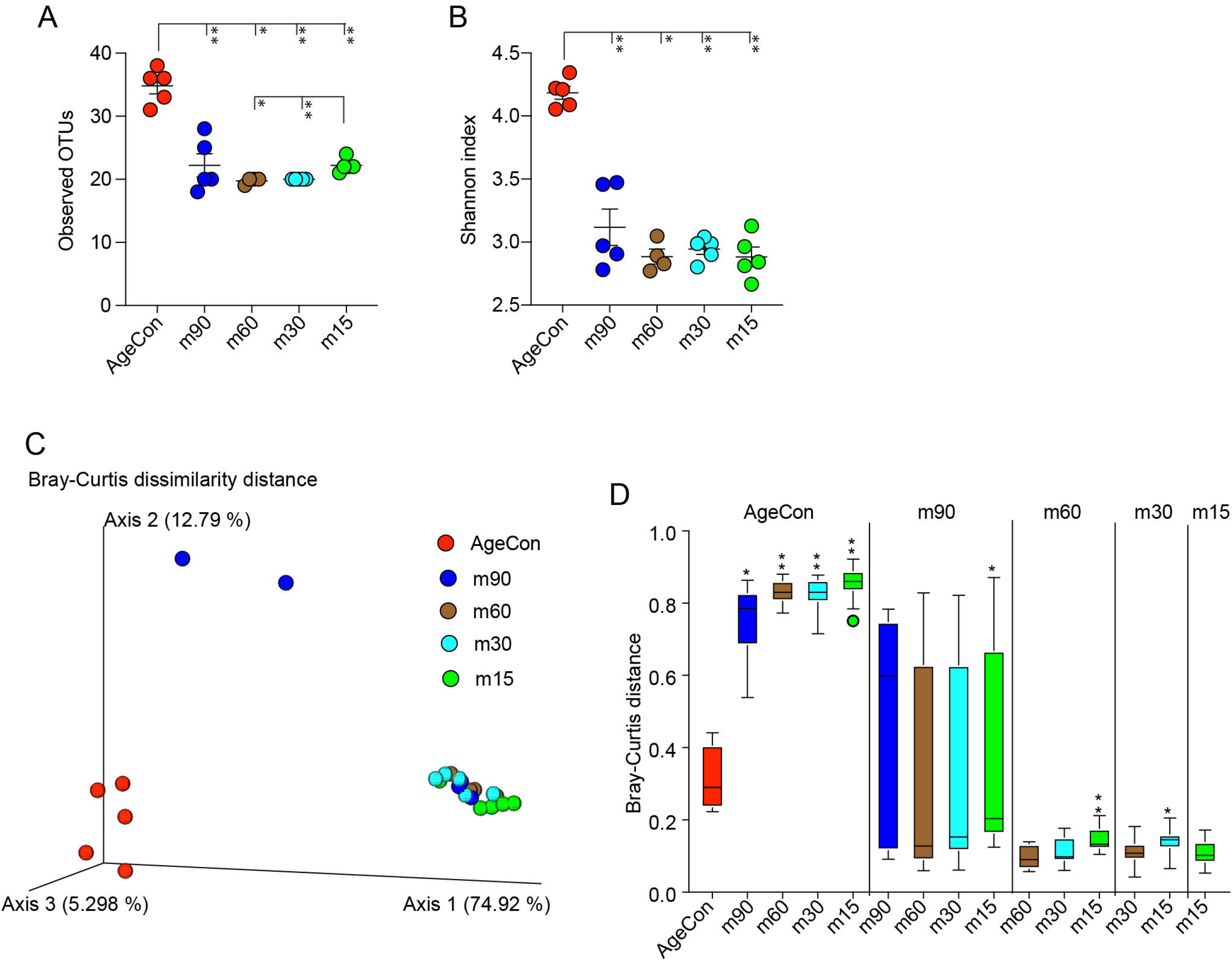
Gut microbiota are not able to recover up to 3 months following vancomycin exposure. Fecal pellets were collected on day 0 prior to *P. yoelii* infection (from Figure 3A) to analyze gut microbiota composition. (**A**) Alpha diversity estimated using observed OTUs. (**B**) Alpha diversity estimated using Shannon index. (**C**) PCoA plot shows beta diversity measured with Bray-Curtis dissimilarity distance. (**D**) Bray-Curtis dissimilarity distance comparisons between groups. Box end shows lower and upper quartile and horizontal line inside box is median. Y-axis shows Bray-Curtis dissimilarity distance of groups on X axis to groups on the top of vertical columns. Data are means ± SEM. (**A** and **B**) Pairwise Kruskal-Wallis test. (**D**) Pairwise PERMANOVA with 999 permutations. Significance level is compared between group at the top of column to group on X axis. Gut microbiota was analyzed from only one experiment of two independent experiments. * = *p* < 0.05, ** = *p* < 0.01.

### Gut bacteria provide continuous cues that dynamically modulate the severity of malaria in mice

The data indicate that gut bacteria present at the time of a *Plasmodium* infection shape the severity of infection. Yet, it remains unclear if the pre-infection bacteria composition imprints a scripted immune response that is fixed, versus a dynamic interaction where gut bacteria provide continuous signals to the host immune system following *Plasmodium* infection that allows for changes in the adaptive immune response and outcome of infection. Two separate approaches, using either CR or Tac mice, were used to test these two possibilities. First, CR mice were treated with vancomycin drinking water for various lengths of time beginning before *P. yoelii* infection, beginning on the day of *P. yoelii* infection, or 7-days after *P. yoelii* infection (Fig. 5A). Consistent with the prior data, CR mice treated with vancomycin at any time before *P. yoelii* infection resulted in reduced *P. yoelii* parasite burden (Fig. 5B-C). Vancomycin treatment starting on the day of *P. yoelii* infection also dramatically reduced parasite burden (Fig. 5B-C). Strikingly, starting vancomycin treatment 7 days post-infection, where parasitemia had already reached ~10%, prevented any further increase in parasitemia, resulting in accelerated parasite clearance and a reduction in parasite burden (Fig. 5B-C).

**Figure 5.**
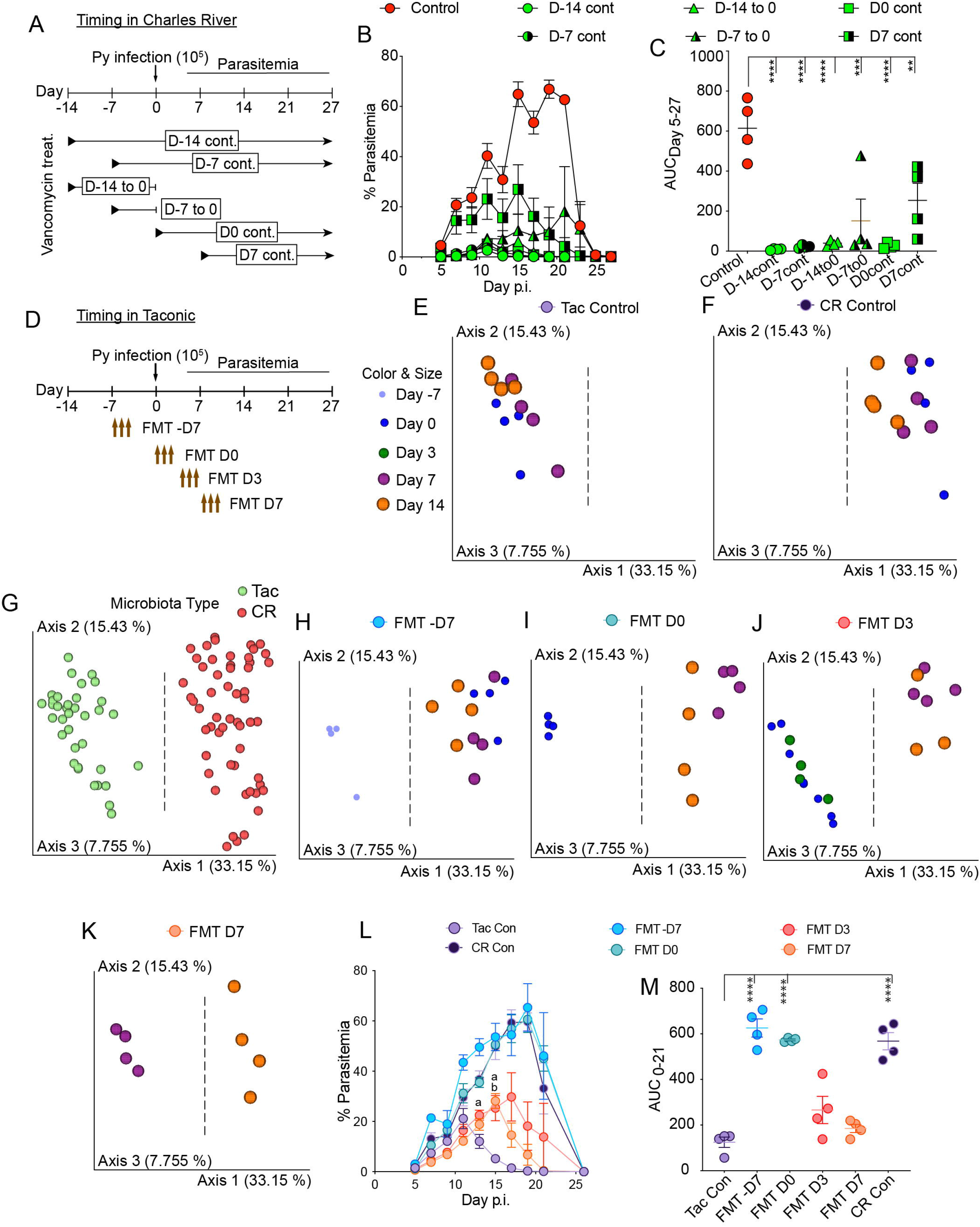
Gut microbiota dynamically modulates *Plasmodium* parasite burden in susceptible and resistant mice. (**A**) C57BL/6 mice from CR mice were treated with vancomycin prior to, at, and after *P. yoelii* infection. N = 4 mice/group. (**B** and **C**) Parasitemia and AUC of CR treated with vancomycin from A. Data are means ± SEM and representative of two or more experiments. (**A-C**) Control: Regular water; D-14 cont: Continuous vancomycin water from day −14 till the end of infection; D-7 cont: Continuous vancomycin water from day −7 till the end of infection; D-14 to 0: Vancomycin water from day −14 to day 0; D-7 to 0: Vancomycin water from day −7 to day 0; D0 cont: Continuous vancomycin water from day 0 till the end of infection and D7 cont: Continuous vancomycin water from day 7 post infection till the end of infection. (**D**) C57BL/6 mice from Tac were gavaged with 3 consecutive cecal contents (FMTs) from CR prior to, at, and after *P. yoelii* infection. N = 4 mice/group. (**E-K**) Gut microbiota analysis using MVRSION approach of Tac and CR mice receiving FMTs were collected at respective days. PCoA plots show beta diversity analysis using Bray-Curtis dissimilarity distance. Vertical dashed line clearly marks the clustering of Tac type to the left and CR type to the right. (Land M) Parasitemia and AUC of Tac mice receiving FMTs from CR from D. Data are means ± SEM and representative of two experiments. (**D-M**) FMT – D7: FMT on day −7, −6, and −5; FMT D0: FMT on day 0, 1, and 2; FMT D3: FMT on day 3, 4, and 5 and FMT D7: FMT on day 7, 6, and 8. (**C** and **M**) one-way ANOVA with Tukey’s multiple comparisons test. Only comparisons with control are shown. ** = *p* < 0.01, *** = *p* < 0.001, and **** = *p* < 0.0001. (**L**) two-way ANOVA with Dunnett’s multiple comparisons test. a = Tac Con versus FMT D3; *p* < 0.05; b = Tac Con versus FMT D7; *p* = 0.002.

In the second approach, Tac mice received CR ceca content transplants for three consecutive days beginning 7 days before *P. yoelii* infection, beginning on the day of *P. yoelii* infection, beginning 3 days after *P. yoelii* infection, or beginning 7 days after *P. yoelii* infection (Fig. 5D). Bacteria community analysis by MVRSION revealed that control Tac mice maintained a differential bacteria community compared to control CR mice through day 14 p.i. (Fig. 5E-F), and that when stool bacteria communities were compared in all samples there were distinct “Tac” and “CR” microbiota types (Fig. 5G). Analysis of each group of Tac mice that received CR FMTs revealed that after the FMT the bacteria communities transitioned to a “CR” microbiota type (i.e., transition from left to right side of dashed line; Fig. 5 H-K) with the FMT D3 group demonstrating this transition occurred within 4 days (comparison of day 3 to day 7 samples; Fig. 5J). As with CR mice, manipulating gut microbiota in Tac mice before or on the day of *P. yoelii* infection resulted in a profound change (i.e., increase in Tac mice) in the parasite burden compared to control Tac mice (Fig. 5L-M). Importantly, Tac mice that received CR cecal content transplants after the *P. yoelii* infection also showed a shift in the parasitemia kinetics (Fig. 5L-M). Collectively, these data demonstrate that gut bacteria provide continuous cues that dynamically modulate the severity of malaria in susceptible and resistant mice.

### Signals from gut bacteria dynamically regulate germinal center reactions and parasite burden

The adaptive immune response, in particular germinal center (GC) reactions and *Plasmodium-specific* antibodies, are required for control of *Plasmodium* blood stage infections (del Portillo et al., 2012). The data in Figure 5 led to the hypothesis that GCs are malleable to the continuous cues provided by gut bacteria, which ultimately determines parasite burden. To test this hypothesis, CR mice were infected with *P. yoelii* (Fig. 6A) and left untreated (noVan), treated with vancomycin beginning on day 0 (d0Van), or treated with vancomycin beginning on day 7 p.i. (d7Van). Each of these groups of mice were also treated with an anti-CD40L monoclonal antibody (clone MR1) that disrupts GC reactions (Foy et al., 1993; Noelle et al., 1992), or with an isotype control antibody (IgG) beginning day 7 p.i. (Fig. 6A). If vancomycin-induced changes in gut bacteria affect *P. yoelii* parasite burden independent of GC reactions, then disrupting GCs would have minimal effect on parasite burden in the vancomycin treated mice. Fecal pellets were collected on day 10 p.i. and MVRSION analysis demonstrated that vancomycin treatment changed the bacterial community (Fig. 6B), with only the d0Van group showing a difference in Bray-Curtis dissimilarity distance between the IgG and MR1 mice within any of the three vancomycin treatment groups (Fig. 6C).

**Figure 6.**
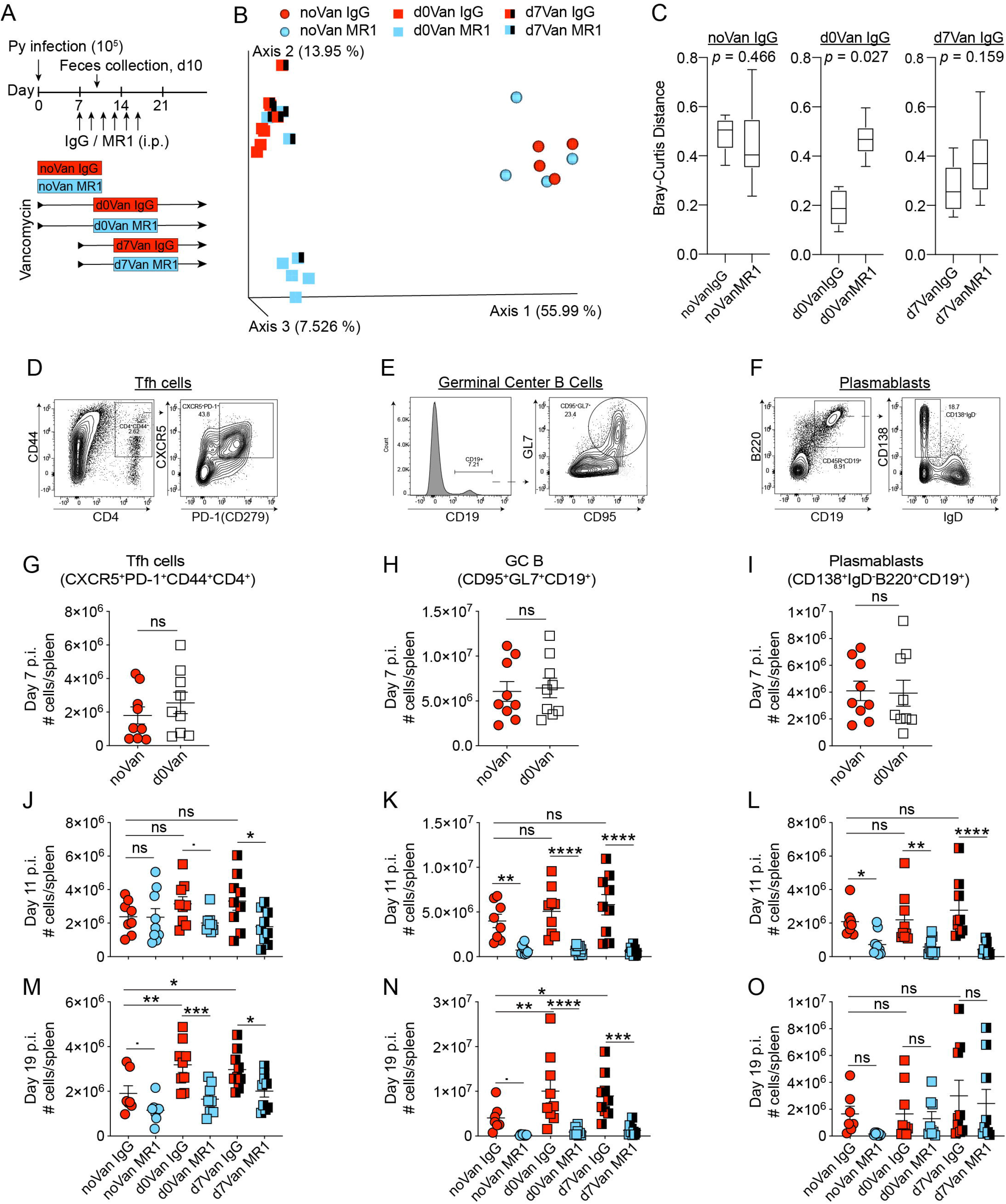
Blocking CD40-CD40L interactions impairs augmented germinal center reactions observed in vancomycin-treated mice. (**A**) C57BL/6 mice from CR mice were continuously treated with vancomycin from day 0 and day 7 post-infection. Mice received either 6 doses of isotype control (IgG) or MR1 intraperitoneally (IP) on alternate days from day 7 to day 17 post-infection. (**B**) PCoA plot shows beta diversity using Bray-Curtis dissimilarity of fecal pellets collected at day 10 p.i. (**C**) Bray-Curtis distance comparison between the groups. Box end shows lower and upper quartile and horizontal line inside box is median. Y-axis shows Bray-Curtis dissimilarity distance of groups on X-axis to groups on the top of vertical columns. (**D-F**) Gating strategy for the indicated cell subsets. Events in the left gate of each cell populations are total splenocytes (i.e., singlets gated from a side scatter and forward scatter plot). (**G-O**) Spleens were harvested on the indicated days for cellular analysis. (**G, J**, and **M**) Tfh cells (CXCR5^+^PD-1^+^CD44^+^CD4^+^) on day 7, 11, and 19 p.i. respectively. (**H, K**, and **N**) GC B cells (CD95^+^GL7^+^CD19^+^) on day 7, 11, and 19 p.i. respectively. (**I, L**, and **O**) Plasmablasts (CD138^+^lgD^-^B220^+^CD19^+^) on day 7, 11, and 19 p.i. respectively. Data are means ± SEM. (**C**) Pairwise PERMANOVA with 999 permutations. Representative of two independent experiments with similar results. N = 3 mice/group. (**G-O**) Cumulative results from three experiments. (**G-I**) T-test. (**J-O**) One-way ANOVA with Sidak correction for selected pairs. noVan IgG: no vancomycin with IgG; noVan MR1: no vancomycin with MR1; d0Van IgG: vancomycin from day 0 and IgG; d0Van MR1: vancomycin from day 0 and MR1; d7Van IgG: vancomycin from day 7 and IgG; and d7Van MR1: vancomycin from day 7 and MR1. · < 0.1, * = *p* < 0.05, ** = *p* < 0.01, ***= *p* < 0.001, and **** = *p* < 0.0001.

Within GC reactions, CD4+ follicular helper T cells (Tfh cells) provide necessary signals to GC B cells to support somatic hypermutation and affinity maturation of antibodies (Victora and Nussenzweig, 2012). GC B cells exit the GC as plasma cells, which produce large quantities of antibodies, or memory B cells. In contrast to prior studies that demonstrated antibiotic treatment impaired adaptive immunity (Abt et al., 2012; Hagan et al., 2019; Ichinohe et al., 2011; Lynn et al., 2018; Oh et al., 2014) vancomycin treatment beginning on the day of *P. yoelii* infection had no effect (noVan versus d0Van) on the number of splenic Tfh cells (Fig. 6D and G), GC B cells (Fig. 6E and H) or plasmablasts (Fig. 6F and I) on day 7 p.i. Additionally, vancomycin treatment beginning on the day of *P. yoelii* infection or day 7 p.i. (noVan IgG versus d0Van IgG or d7Van IgG, respectively) did not result in fewer numbers of splenic Tfh cells (Fig. 6J), GC B cells (Fig. 6K) or plasmablasts (Fig. 6L) on day 11 p.i. Intriguingly, over the course of infection, vancomycin treatment (d0Van IgG or d7Van IgG compared to noVan IgG) resulted in increased numbers of Tfh cells and GC B cells with a more modest trend also observed within the plasmablasts (Fig. 6M-O and Fig. S10). Collectively, the data demonstrate that the effect of antibiotics is not necessarily detrimental to host immunity, but in certain contexts may support a more robust adaptive immune response.

To assess the efficiency of MR1 treatment that began day 7 p.i. at blocking GC reactions, the number of Tfh cells, GC B cells, and plasmablasts were also quantified in mice that received MR1. At day 11 p.i., MR1 treatment had little effect on the number of Tfh cells except between the d7Van IgG and d7Van MR1 treated mice (Fig. 6J). In contrast, MR1 treatment resulted in significant decreases in both GC B cells (Fig. 6K) and plasmablasts (Fig. 6L) in all three groups of mice (noVan, d0Van, and d7Van). On day 19 p.i., MR1 treatment, compared to IgG, resulted in fewer Tfh cells (Fig. 6M) and a sustained loss in GC B cells (Fig. 6N) between the noVan, d0Van, and d7Van groups of mice, while no significant effect on numbers of plasmablasts (Fig. 6O). Collectively, these data demonstrate that vancomycin treatment, even up to one-week post-P. *yoelii* infection, rapidly changes gut bacteria populations, increases numbers of GC B cells, and that MR1 treatment one-week post infection was sufficient at ablating the elevated GC responses observed in vancomycin treated mice.

Finally, parasite burden was analyzed to determine if signals from gut bacteria impact the severity of malaria through dynamic modulation of GC reactions in CR mice (Fig. 7A). Consistent with GCs required for control of *Plasmodium*-infected red blood cells, MR1 treatment in the noVan group resulted in a continuous increase in parasite burden through the end of MR1 treatment followed by a pronounced delay in clearance (Fig. 7B-C). Mice treated with vancomycin on day 0 showed the expected low parasite burden, however disruption of GC reactions in these mice beginning on day 7 p.i., even though parasite burden was already quite low compared to the noVan mice, resulted in a significant increase in parasite burden (Fig. 7D-E). Moreover, MR1 treatment also ablated the low parasite burden that was observed in d7Van IgG treated mice (Fig. 7F-G). These data indicate that gut bacteria dynamically interact with GC reactions to impact *Plasmodium* infection, and that certain microbiota composition can lead to enhanced host protection.

**Figure 7.**
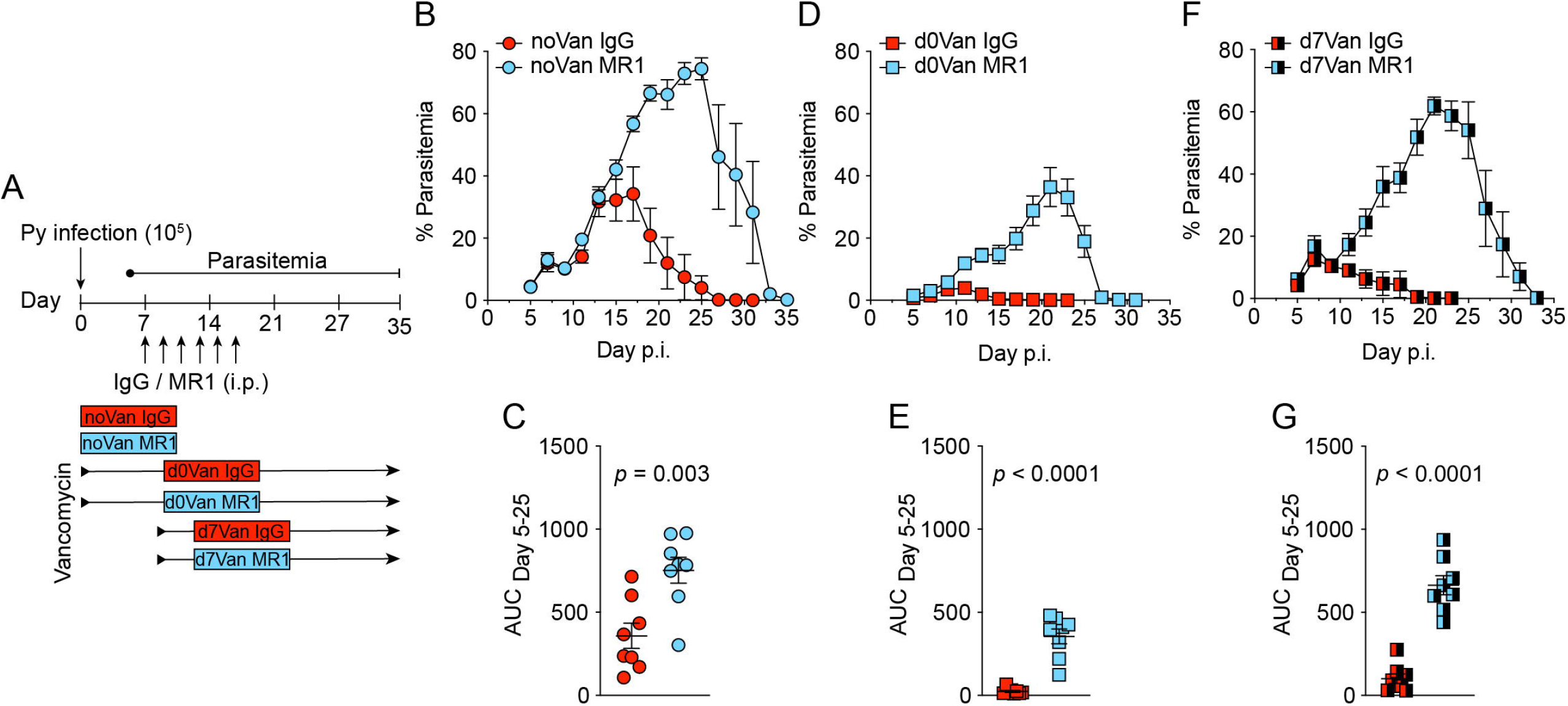
Gut bacteria dynamically interact with germinal center reactions to modulate *Plasmodium* parasite burden. (**A**) Experiment design. (**B, D**, and **F**) Parasitemia on the indicated days. (**C, E**, and **G**) Area under curve (AUC) analysis. Data are means ± SEM. (**C, E**, and **G**) T-test. Data are cumulative from two independent experiments. N = 8 mice/group.

## Discussion

These results provide an intriguing observation that gut microbiota modulated before or during active *Plasmodium* infection can alter host protection and disease prognosis. This stems from the malleable nature of GC reactions following *Plasmodium* infection and the ability of GC reactions to respond to cues from gut bacteria, which ultimately affect parasite burden. GCs are anatomical sites within lymph nodes and the spleen where somatic hypermutation-induced affinity maturation of antibodies, clonal B cell expansion and generation of long-lived antibody secreting plasma cells and memory B cells occur to provide protection against infection and reinfection (Victora and Nussenzweig, 2012). As a *Plasmodium* infection is blood-borne, the majority of the immune responses involved in the clearance of the parasite occur in the spleen (del Portillo et al., 2012). GCs form in the spleen of mice by day 6 post-*Plasmodium* infection (Ryg-Cornejo et al., 2016). The development of GC reactions during the first week of infection, through the continuous cues provided by gut bacteria, is quite dynamic and is critical to disease outcome as evidenced by these data. Important questions remain, including what cells of the immune system do gut microbiota continuously signal through to modulate GC reactions and what components of gut bacteria (e.g., proteins, polysaccharides, biochemical metabolites, small RNAs, etc.) regulate GC reactions. Prior work has shown that short-chain fatty acids from gut bacteria can impact GC B cells (Kim et al., 2016). Yet, Tac and CR mice do not show differences in serum short-chain fatty acid levels (Chakravarty et al., 2019), suggesting that these metabolites are not responsible for dynamic modulation of GC reactions during *Plasmodium* infection.

We report here that antibiotic-induced changes in gut bacteria were beneficial to CR mice as evident by the decreased blood stage *Plasmodium* parasite burden. In contrast, numerous studies have shown that oral antibiotics lead to gut bacteria dysbiosis that have detrimental effects on host health. For example, antibiotic-induced gut microbiota perturbation leads to increased inflammation in blood and alters immunity to vaccines in humans (Hagan et al., 2019). A single dose of intraperitoneal clindamycin leads to sustained susceptibility to *Clostridioides difficile* induced diarrhea and colitis in mice due to significant alterations of intestinal microbiota (Buffie et al., 2012). Similarly, antibiotic use in neonates and adults predispose to wide range diseases like diabetes, obesity, inflammatory bowel diseases, asthma, depression, and autism, among others (Zhang and Chen, 2019). In contrast to these scenarios, and in agreement with this report, a cocktail of antibiotics (ampicillin, metronidazole, neomycin, and vancomycin) were advantageous to the host as they prevented motor dysfunction and limited axon damage in a murine model of progressive multiple sclerosis (Mestre et al., 2019). Yet, this cocktail of antibiotics, which eliminates the vast majority of gut bacteria altogether, creates a different scenario than in the present study, in which single antibiotics were used to induce altered bacteria compositions.

Whereas antifungal and antiviral treatments on *P. yoelii* parasitemia were inconsistent or minimal, respectively, there is the possibility that fungi and viruses that contribute to modulating parasitemia were not affected by the treatments used in this study. Thus, further studies are necessary to fully assess the role of these gut microbes in the regulation of *Plasmodium* parasite burden. The present data indicate that gut bacteria are the critical gut microbes that effect parasitemia. Curiously, the effect of gut bacteria on the blood stage parasite burden was not attributed specifically to Gram-positive, Gram-negative, or anaerobic bacteria, as antibiotics that target these different classes of bacteria were capable of decreasing parasite burden in CR mice. Therefore, it appears to be the consortium of gut bacteria that affects parasitemia rather than a single class of bacteria. Importantly, the effects of antibiotic treatment on parasitemia were dependent on baseline gut microbiota composition as different results were observed in Tac and CR mice. These observations are consistent with a prior report in which mice colonized with unrelated human stool samples, resulting in different baseline gut microbiota communities, had altered colon transcriptomes following amoxicillin-clavulanate treatment (Lavelle et al., 2019).

One of the striking observations in this study was that gut bacteria composition remain altered for up to 3-months following a 14-day vancomycin treatment in drinking water. Prior studies have shown that mice treated with broad spectrum antibiotics (ciprofloxacin and metronidazole) for 2 weeks in drinking water with 4 weeks of spontaneous recovery had partially restored gut microbiota to baseline level (Suez et al., 2018). Consistently, gut microbiota of healthy human adults returned to near-baseline composition within 1.5 months following 4-day oral antibiotic (meropenem, gentamicin, and vancomycin) treatment (Palleja et al., 2018). In contrast, in a study in which healthy human adults were treated with broad-spectrum antibiotics (ciprofloxacin, vancomycin, and metronidazole) for 7 days it took 8-31 months for gut bacteria to return to pre-treatment levels, and even then the gut microbiota community composition was slightly altered from baseline (Haak et al., 2018). These data suggest that the recovery of gut microbiota likely depends on type of antibiotics, drug dose regimen, and host. Similarly, other factors may affect the recovery such as host diet, community context, environmental reservoirs, post-antibiotic environment factors like probiotic intake and autologous FMT (Ng et al., 2019; Suez et al., 2018).

As vancomycin treatment beginning the day of *P. yoelii* infection resulted in low parasitemia (Fig. 5B and Fig. 7B,D), in spite of no increase in GC-associated cell numbers through day 11 p.i. (Fig. 6G-L), this suggests that vancomycin-induced decreases in parasitemia early during infection may be the result of changes in the quantity or quality of the *P. yoelii-*induced antibodies. Alternatively, whereas GC are required for the ultimate clearance of infected red blood cells (Figueiredo et al., 2017; Guthmiller et al., 2017; Obeng-Adjei et al., 2015; Pérez-Mazliah et al., 2015; Pérez-Mazliah et al., 2017; Ryg-Cornejo et al., 2016), it is possible that the reduction in parasitemia early during infection following vancomycin treatment that began the day of *P. yoelii* infection may have occurred through augmenting other aspects of the host immune response. Given the complexity of the innate and adaptive immune response to *Plasmodium*, assessing whether and how vancomycin-induced changes in gut microbiota modulate additional components of the immune response to *Plasmodium* will require future investigation.

It was recently shown that Kenyan children having two episodes of malaria have different gut microbiota compositions than children having only one episode (Mandal et al., 2018). This observation further supports the possibility that gut bacteria in humans may impact the severity or outcome of *Plasmodium* infection. Excitingly, for the first time, to our knowledge, we have reported that gut bacteria composition in Uganda children differs between asymptomatic *P. falciparum* infection and severe malarial anemia. The observation that stool bacteria populations in Ugandan children with SMA collected at varying times post-enrollment and treatment were not different is also consistent with the prior study in Kenya showing that malaria episodes did not alter stool bacteria populations (Mandal et al., 2018). Collectively, these observations provide tantalizing support that the gut microbiome may shape the severity of malaria in children. Nevertheless, future studies incorporating longitudinal analysis (i.e., collection human stool samples before and through malaria transmission seasons) are necessary to further demonstrate the potential for baseline gut microbiota to modulate the severity of malaria in children. Finally, shotgun metagenomics, culturomics, clinical trials, etc. need to be performed to fully decipher links between specific gut bacteria and severity of malaria in humans (Mukherjee et al., 2019). As the decline in the number of malaria deaths has largely plateaued in recent years, and since the gut microbiome has significant impacts on host health and is amendable to change via diet, prebiotics and/or probiotics, gut microbiota-based therapeutics may represent novel approaches to prevent severe malaria and deaths.

This report demonstrates that gut bacteria in mice continuously interact with gut-distal germinal center reactions during an ongoing extra-gastrointestinal tract infection to shape the severity of disease. Importantly, gut bacteria composition was also shown to associate with the severity of malaria in humans, suggesting that human gut microbiota may represent a novel target to ameliorate severe malaria. Finally, these data demonstrate that antibiotic-induced changes in gut bacteria do not uniformly impair systemic adaptive immunity, but can in in some cases result in elevated immune responses. These results highlight the need for ongoing research to investigate the diverse interactions between gut microbiota and host immunity.

## Materials and Methods

### Animals and housing

Genetically similar 6-8 weeks old female C57BL/6N mice were obtained from Taconic Biosciences (Isolated Barrier Unit^™^ IBU050401C; Hudson, NY) and Charles River Laboratories (barrier room R01; Wilmington, MD) and housed conventionally in a specific pathogen-free (SPF) facility. Mice were kept on NIH-31 Modified Open Formula Mouse/Rat Irradiated diet (Harlan 7913) (Harlan, Indianapolis, IN) and non-acidified autoclaved reverse osmosis water ad libitum upon arrival. Mice were acclimatized for a minimum of one week prior to performing experiments. Mice were kept on a 12-hour light (6 AM – 6 PM) and 12-hour dark (6 PM – 6 AM) cycle. Animal handling and experiment protocols were approved by the University of Louisville and Indiana University Institutional Animal Care and Use Committees (IACUC).

### *Plasmodium* infection and parasitemia analysis

Donor mice were injected intravenously (IV) with thawed 10^5^ infected red blood cells (iRBCs) with either *Plasmodium yoelii* 17XNL or *Plasmodium berghei* ANKA. Blood was collected on day 5 post infection (p.i.) via retro-orbital bleed. The blood was counted for iRBCs with Giemsa stain and total RBCs (using Hemocytometer) and diluted in 0.9% saline (Teknova, Hollister, CA) at a concentration of 10^5^ iRBCs / 200 μl. Experimental mice were infected with 10^5^ iRBCs IV unless indicated.

For parasitemia, approximately 5 μl whole blood was collected from tail snip in cold 100 μl 1X PBS in a 96 well plate on ice. The cells were fixed in 0.00625% glutaraldehyde, stained with conjugated antibodies, and subjected flow-cytometry analysis for evaluation of percent parasitemia. The conjugated antibodies for staining panel were CD45.2-APC (clone 104; Biolegend, San Diego, CA), Ter 119-APC/Cy7 (clone TER-119; Biolegend, San Diego, CA), dihydroethidium (MilliporeSigma, St. Louis, MO), and Hechst 33342 (MilliporeSigma, St. Louis, MO). Forward and side scatter singlets were gated on Ter119^+^CD45.2^-^ for RBC. RBCs were gated on Hoechst^+^Dihydroethidium^+^ to calculate the percent of iRBC (% Parasitemia). Parasitemia was tacked every other day beginning day 5 to clearance of parasite unless indicated.

### Cecal microbiota transplantation

Cecal donor mice were anesthetized with isoflurane and cervical dislocation was performed inside a laminar flow hood. Mice were laid on the back and ventral side were doused with 70% ethanol. Ceca were aseptically removed, and contents were squeezed on Petri plate with 2 ml saline. Cecal content was mixed with 1ml syringe plunger. The diluted ceca contents were drawn into a 1ml syringe and 200 μl content were gavaged per mouse. A sterile gavage needle and syringe was changed between each group.

### Antimicrobials

Mice were treated with antimicrobials in drinking water. Antibiotics used were ampicillin (0.5 mg/ml) (Sigma Aldrich; St. Louis, MO), gentamicin sulfate (0.5 mg/ml) (Corning; Manassas, VA), metronidazole (0.5 mg/ml) (Spectrum Chemical, Gardena, CA), neomycin sulfate (0.5 mg/ml) (EMD Millipore; Billerica, MA), and vancomycin (0.25 151 mg/ml) (Amnesco; Solon, OH). Antifungals were fluconazole (0.5 mg/ml) (Sigma-Aldrich), Amphotericin B (0.1 mg/ml) (Sigma-Aldrich), and 5-flurocytosine (1 mg/ml) (Sigma-Aldrich). Antivirals used were ribavirin (0.13 mg/ml), lamivudine (0.05 mg/ml) (GlaxoSmithKline), and Acyclovir (0.1 mg/ml) (GlaxoSmithKline). All of the antimicrobials dissolved in non-acidified autoclaved reverse osmosis drinking water except Amphotericin B (VWR Life Science, CA) which was solubilized in DMSO (VWR Life Science, CA). Antimicrobial water was changed every week.

### Stool and blood collection from Ugandan children

The study was reviewed and approved by the Makerere University School of Medicine Research and Ethics Committee, Indiana University School of Medicine Institutional Review Board, and the Ugandan National Council for Science and Technology. Written and informed consent was obtained from parents or guardians prior to enrolment.

Children between the ages of 0.5 to 4 years old, with 5 most common clinical manifestations of severe malaria (cerebral malaria, CM; respiratory distress, RD; severe malarial anemia, SMA; malaria with complicated seizures, M/S; and prostration) were enrolled in a prospective longitudinal cohort study at two sites: 1) the Pediatric Acute Care Unit at Mulago National Referral Hospital in Kampala, Uganda, and 2) the Pediatric Emergency Ward at the Jinja Regional Referral Hospital in Jinja, Uganda. Children from the same neighborhoods without active illness or fever during enrollment were enrolled in the healthy community control (CC) group. SMA was defined as *P. falciparum* smear or RDT positive and serum hemoglobin level ≤ 5 g/dL All study participants (SMA or CC) were tested for parasitemia by microscopy or RDT and those determined to be parasite positive were treated immediately, according to the Ugandan Ministry of Health treatment guideline at the time of diagnosis. Stool samples from 40 children with SMA, 7 Pf Pos, and 28 Pf Neg children from the parent study, with parasite prevalence confirmed by microscopy, were included in this gut microbiome study. Stool samples were collected various days after enrollment from SMA children (Supplemental Table 1) and during enrollment from CC (Pf Pos + Pf Neg) children and frozen immediately at −80°C. Stool samples were kept frozen until DNA extraction. For blood parameters and parasite detection, peripheral blood was collected by venipuncture on hospital admission from SMA and as an outpatient from CC. A complete blood count (CBC) with differential and reticulocyte count and a blood smear for malaria parasite count were conducted. HIV testing was done on any participants whose parents/guardians did not opt out of testing. There was only two HIV positive participant, which was in the SMA group. No power outage or other situations were noted during samples handling and processing to affect the stability of samples.

### Gut Microbiota Analysis

DNA from feces was extracted using QIAamp PowerFecal DNA kit (QIAGEN, Germantown, MD) according to the manufacturer’s instructions. Extracted DNA samples were shipped overnight on ice packs to the Genome Technology Access Center (GTAC; Washington University, St. Louis, MO) for 16S rRNA sequencing unless indicated. rRNA were sequenced using an approach, Multiple 16S Variable Region Species-level Identification (MVRSION), that can sequence all the 9 hypervariable regions of 16S rRNA gene with 12 primer pairs (Schriefer et al., 2018). Observed operational taxonomic unit (OTU) table were constructed by GTAC. OTU table was imported inside QIIME2 for core-diversity and statistical analysis.

For analysis of bacteria communities in the reciprocal ceca transplants (S Figure 1), DNA samples were shipped overnight on ice packs to the Integrated Microbiome Resource within the Centre for Comparative Genomics and Evolutionary Bioinformatics at Dalhousie University (IMR-CGEB, Halifax, NS, Canada). V6-V8 regions of 16S rRNA were sequenced and analyzed as described previously (Denny et al., 2019). Briefly, paired end sequences were stitched together, low quality reads and chimera were removed, and OTU table was constructed using open reference method using SortMeRNA v2.0.

Sample metadata values were predicted with supervised machine learning algorithm using random forest with q2-sample-classifier plugin using QIIME2. One-third of the samples were used as test set data. 15 and 20 K-fold cross-validation were performed in mice and human study, respectively.

### Spleen immune cell analysis

RP10 media: RPMI 1640 media (Thermo Fisher Scientific Inc., Waltham, MA) supplemented with 10% fetal bovine serum (FBS) (Atlanta Biologicals, Inc., Lawrenceville, GA), 1.19 mg/ml HEPES (Thermo Fisher Scientific Inc., Waltham, MA), 0.2 mg/ml L-glutamine (Research Products International Corp., Mt. Prospect, IL), 0.05 units/ml & 0.05 mg/ml penicillin/streptomycin (Invitrogen, Grand Island, NY), 0.05 mg/ml gentamicin sulfate (Invitrogen, Grand Island, NY), and 0.05 μM 2-Mercaptoethanol (Thermo Fisher Scientific Inc., Waltham, MA). Spleens on indicated days were harvested in RP10 media and smashed on a wire sheath with flat side of 10 ml syringe plunger to make single cell suspension. Single cell suspensions were washed once with RP10 and treated with ACK lysis buffer (NH_4_Cl −150 mM, KHCO_3_–1 mM, and Na_2_EDTA–0.1 mM in distilled water) to lyse red blood cells. Single cell suspensions were counted and resuspended in RP10 at 2X10^7^ cells/ml.

Cells (2-10^6^) were stained in FACS buffer (1X PBS, 1%FBS, and 0.02% sodium azide) for 30 min at 4C with the following fluorescence-conjugated antibodies (CD45.2, clone 104; CD4, clone RM4-5; CD8, clone 53-6.7; CD19, clone 6D5; Ter119, clone Ter-119; CD11a, clone M17/4; CD49d, clone R1-2; PD-1, clone 29F.1A12, CD95, clone Jo2; GL7, clone GL7; biotion-CXCR5, clone 2G8; CD44, clone IM7; B220, clone RA3-6B2; CD138, clone 281-2; IgD, clone 11-26c.2a) purchased from Biolegend (San Diego, AC) and BD Biosciences (San Diego, CA). For CXCR5 staining, cells were first stained with biotinylated CXCR5 at room temperature for 30 min and then stained with fluorescence-conjugated streptavidin. Cells were fixed and permeabilized with fixation buffer (Biolegend, San Diego, CA) after staining. Cells were acquired with a BD LSRFortessa (BD Biosciences, San Jose, CA). Data were analyzed by FlowJo software (Tree Star, Ashland, OR)

### Statistical analysis

Statistical analyses were performed using GraphPad Prism 6 software (GraphPad), QIIME2, and R packages. Specific statistical tests are described in the figure legends. For area under the parasitemia curve (AUC) analyses, the trapezoidal rule was used for equation:

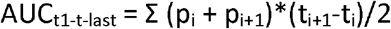

where “p” is percent parasitemia at the designated time point “t” (Méndez et al., 2006).

Significance level of alpha diversity between Pf Pos and SMA (malaria severity) were tested with ANOVA using following model accounting for age (AgeYrs) and sex.

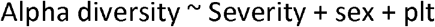

Significance level of beta diversity between Pf Pos and SMA (malaria severity) were tested with “Adonis” function with “bray” method of vegan package implemented in R with following model accounting for age, antibiotics use, percent neutrophil, and weight for height z-score (whz).

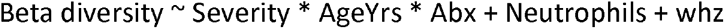

## Data availability

The 16S rRNA gene sequencing datasets generated and analyzed from mice and humans in this study are publicly available and deposited in the NCBI Sequence Read Archive under the BioProject IDs PRJNA643559 and PRJNA642859 respectively.

## Acknowledgements

The authors thank Dr. Martin Richer for critical review of this manuscript. This work was supported by a grant from the National Institute of Allergy and Infectious Disease of the National Institutes of Health (1R01AI123486 to N.W.S. and R01NS055349 to C.C.J.) and funds from the University of Louisville to N.W.S. Support provided by the Herman B Wells Center (N.W.S. and C.C.J.) was in part from the Riley Children’s Foundation. The project described was supported by the Indiana University Health – Indiana University School of Medicine Strategic Research Initiative to N.W.S. The Indiana University Melvin and Bren Simon Cancer Center Flow Cytometry Resource Facility is funded in part by NIH, National Cancer Institute grant P30 CA082709, National Institute of Diabetes and Digestive and Kidney Diseases (NIDDK) grant U54 DK106846, and by NIH instrumentation grant 1S10D012270. The content is solely the responsibility of the authors and does not necessarily represent the official views of the National Institutes of Health. *S.D.G*.

## Author Contribution

RKM: Designed and performed experiments; analyzed data; drafted, reviewed and edited manuscript.

JED: Performed experiments, reviewed and edited manuscript.

RN: Design data collection from Uganda Children; reviewed manuscript.

ROO: Design data collection from Uganda Children; reviewed manuscript.

DD: Data collection from Uganda Children; reviewed and edited manuscript

CCJ: Design and data collection from Uganda children; reviewed and edited manuscript.

NWS: Designed experiments; drafted, reviewed and edited manuscript.

## Supplementary Materials

**Figure S1.**
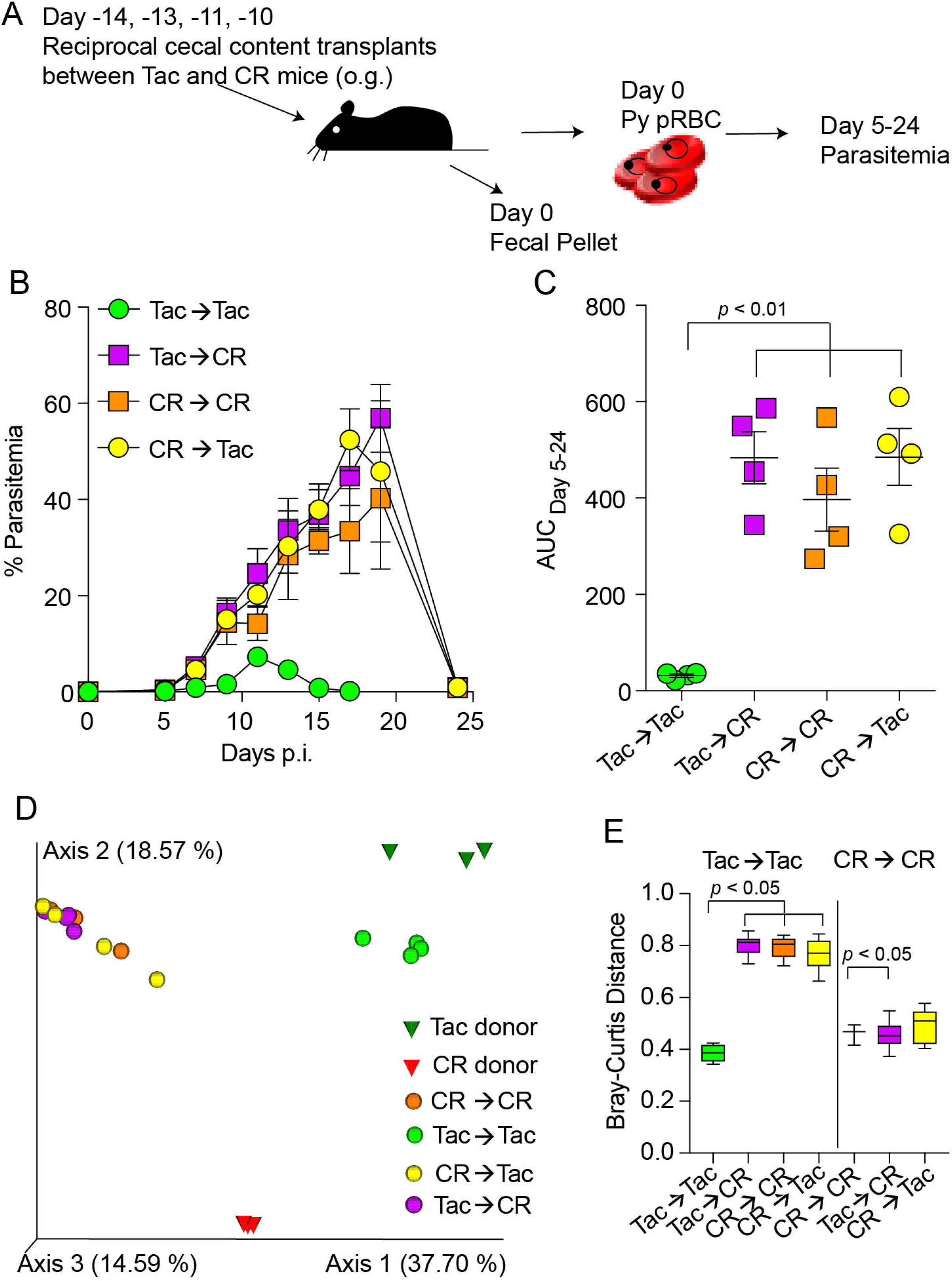
Susceptible gut microbiota are dominant over resistant microbiota. (**A**) Cecal microbiota of C57BL/6 mice from two suppliers, Taconic (Tac) and Charles River (CR), were gavaged reciprocally with cecal contents in saline at days −14, −13, −11, and −10 prior to *P. yoelii* infection; parasitemia was tracked. (**B**) Percentage of RBCs infected with *P. yoelii* (percent parasitemia). N = 4 per group; data are representative of multiple experiments. (**C**) Area under curve analysis (AUC) of (**B**). (**D**) Gut microbiota analysis from mice in (**B**) at day 0 using V6-V8 region of 16S rRNA gene. Principal coordinate analysis showing beta diversity analysis. Cecal contents were sequenced for donors while fecal pellets were sequenced for the recipients. Gut microbiota analysis is from shown experiment. (**E**) Bray-Curtis dissimilarity distance comparison between the groups. Box end shows lower and upper quartile and horizontal line inside box is median. Y-axis shows Bray-Curtis dissimilarity distance of groups on X-axis to groups on the top of vertical columns. Data are means ± SEM. (**C**) One-way ANOVA with Tukey’s multiple comparisons test. (**E**) Pairwise PERMANOVA with 999 permutations.

**Figure S2.**
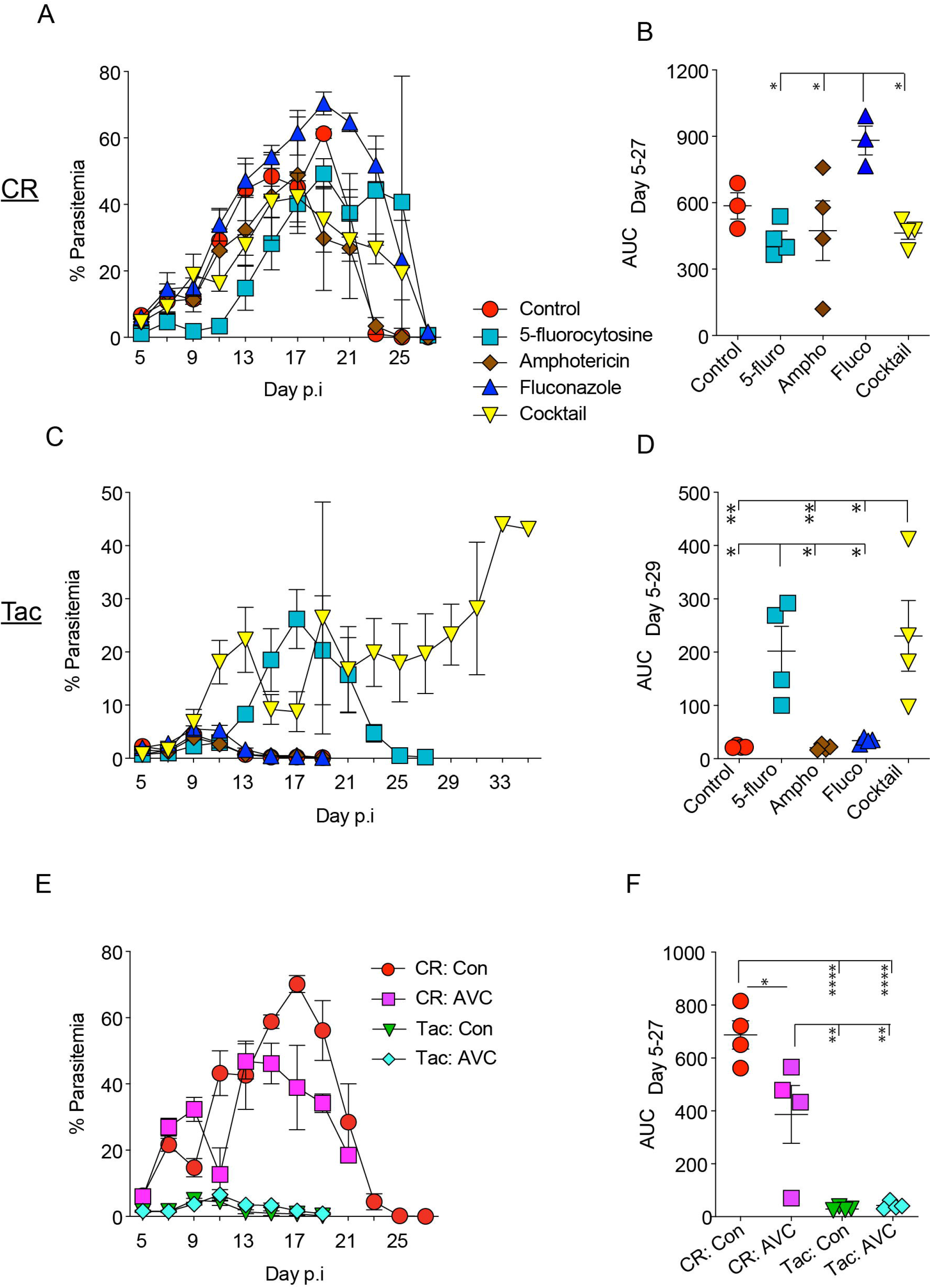
Antifungals and antivirals have a modest impact on severity of malaria. Mice were treated with antifungals and antivirals from 2 weeks prior to infection till the end of infection. Mice were infected with 10^5^ infected *P. yoelii* RBC on day 0. (**A**, and **C**) Parasitemia of CR and Tac mice treated with antifungals respectively. (**B**, and **D**) AUC analysis of CR and Tac mice treated with antifungals respectively. (**E**) Parasitemia of CR and Tac mice treated with antiviral cocktail. (**F**) AUC of CR and Tac mice treated with antiviral cocktail. Data are means ± SEM. (**B, D**, and **F**) Statistical significance were analyzed by one-way ANOVA with Tukey’s multiple comparisons test. Representative of two independent experiments with similar results. N = 4 per group in each experiment. * = *p* < 0.05, ** = *p* < 0.01, *** = *p* < 0.001.

**Figure S3:**
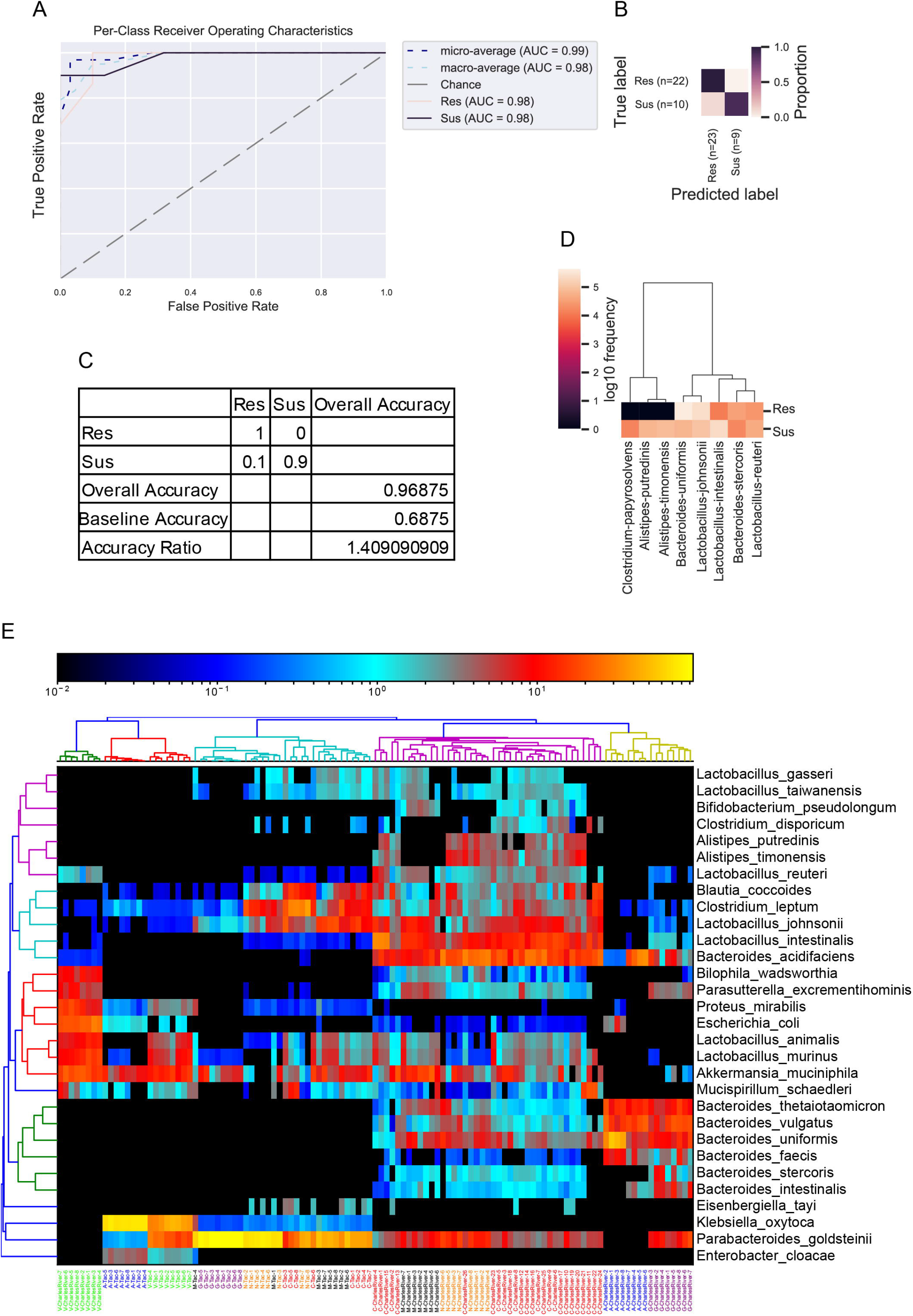
Predicting malaria phenotype in mice using random forest. Groups of CR and Tac mice were categorized as either susceptible malaria phenotype (untreated control CR mice plus CR mice treated with neomycin) or resistant malaria phenotype (CR mice treated with ampicillin, gentamicin, or vancomycin plus untreated control Tac mice or Tac mice treated with one of the five antibiotics). CR mice treated with metronidazole were not included in these analyses because of intestinal absorption and direct inhibitory effect on *Plasmodium*. (**A**) Per-class receiver operating characteristics curve. (**B**) Result of prediction with test set data. (**C**) Accuracy of machine-learning model. (**D**) Bacterial species that are predictive of the susceptible and resistant phenotypes. (**E**) Heatmap shows relative abundance of top 30 abundant species. Column names represent mice treated with antibiotics using different color codes. Hierarchical clustering of samples and features were performed using average linkage method.

**Figure S4:**
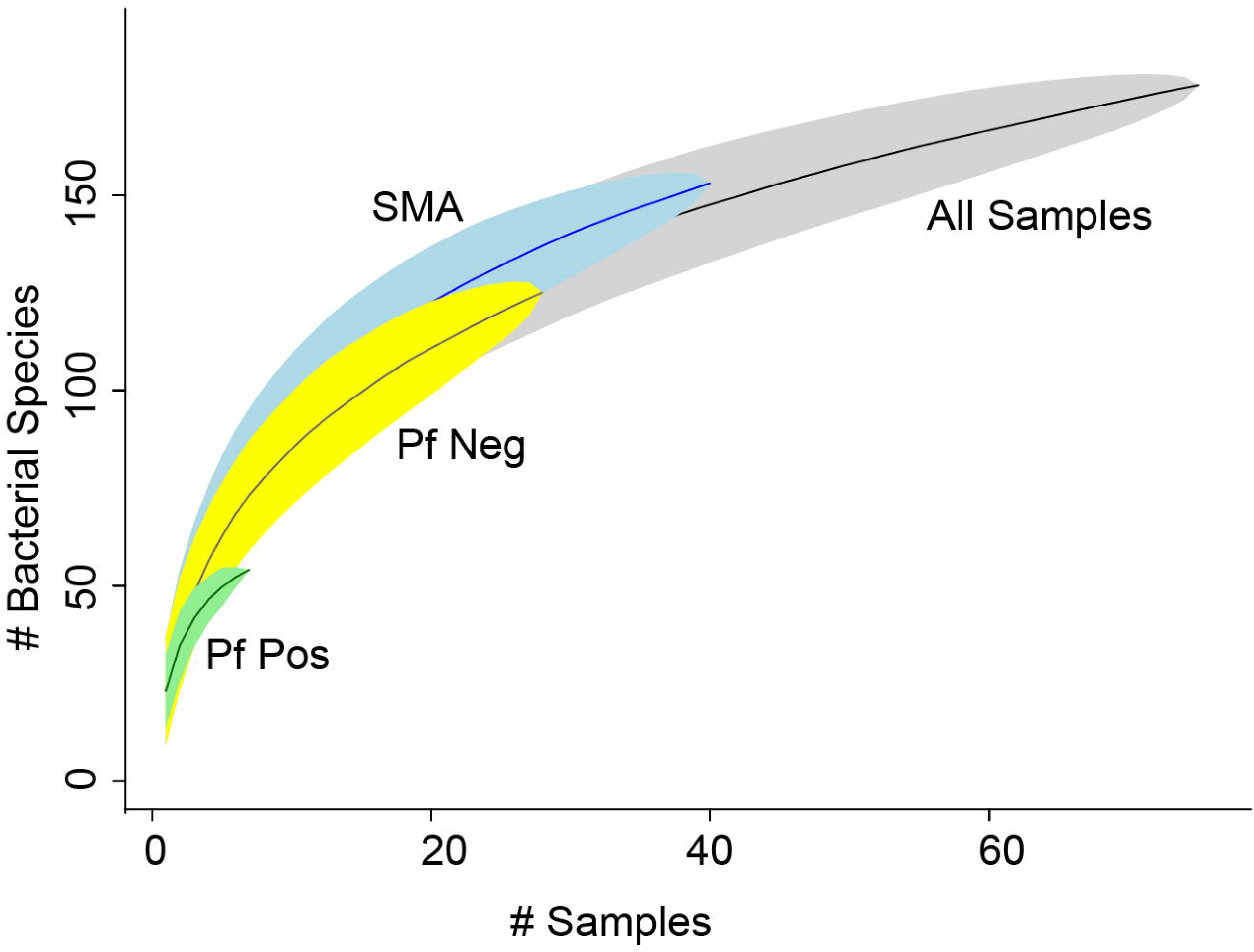
Species accumulation curve shows sufficient number of samples in SMA and Pf pos samples in Uganda children. Shaded area indicates 95% confidence interval. Graph was constructed using *specaccum* function with “random” method using vegan package in R.

**Figure S5.**
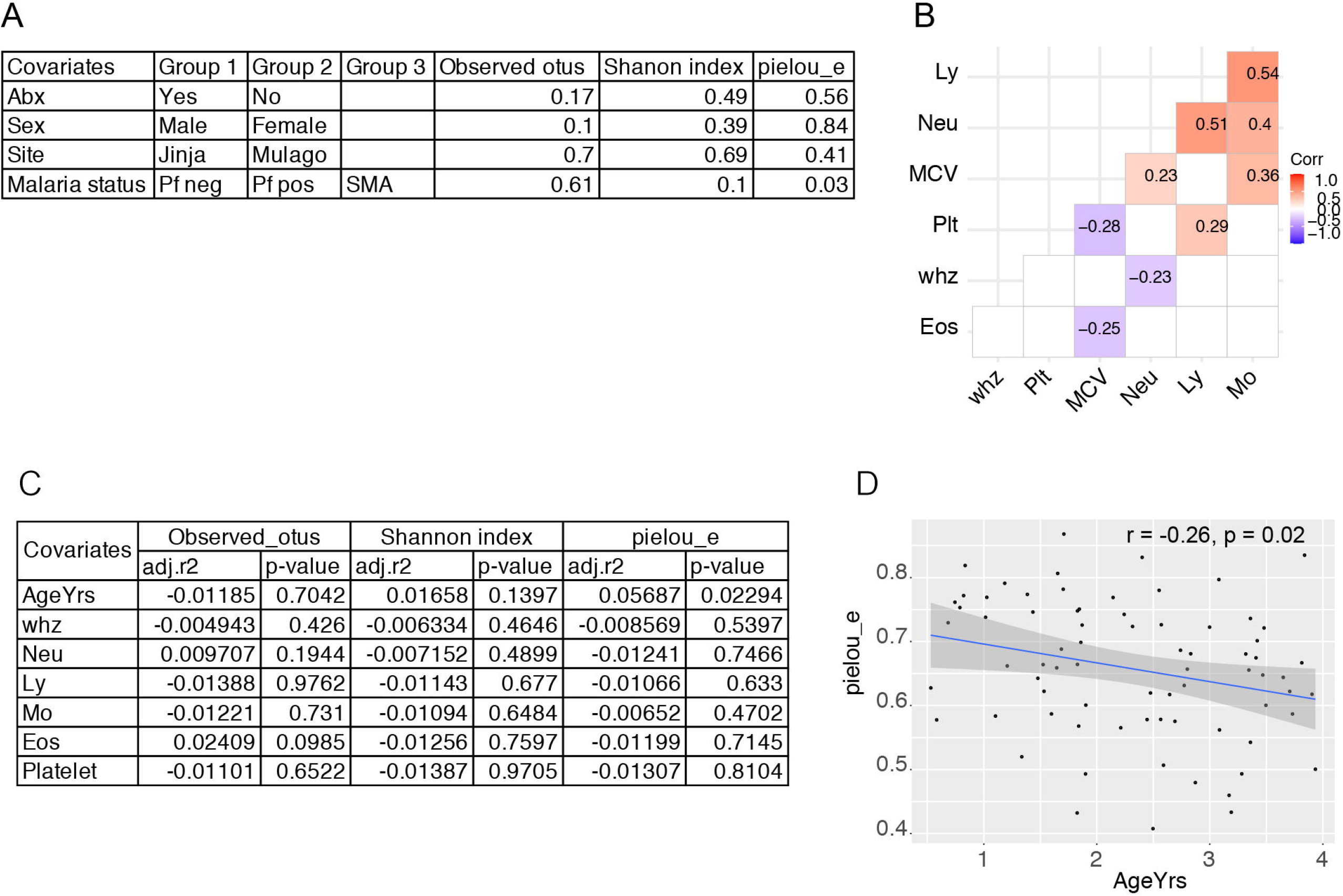
Alpha diversity analysis of gut microbiota in children with different *Plasmodium* status. (**A**) Effect of categorical covariates of interest on alpha diversity. (**B**) Correlation matrix of continuous covariates of interest. Neu, Eos, Ly, Mo, Plt – Absolute count (K/mcl) of blood parameters Neutrophils, Eosinophils, Lymphocytes, Monocytes, and Platelets respectively; MCV – mean corpuscular volume (fL); and whz – weight for height z-score. (**C**) Effect of continuous covariates of interest on alpha diversity measured using regression analysis. (**D**) Correlation of alpha diversity measured using pielou_e and age of participants.

**Figure S6.**
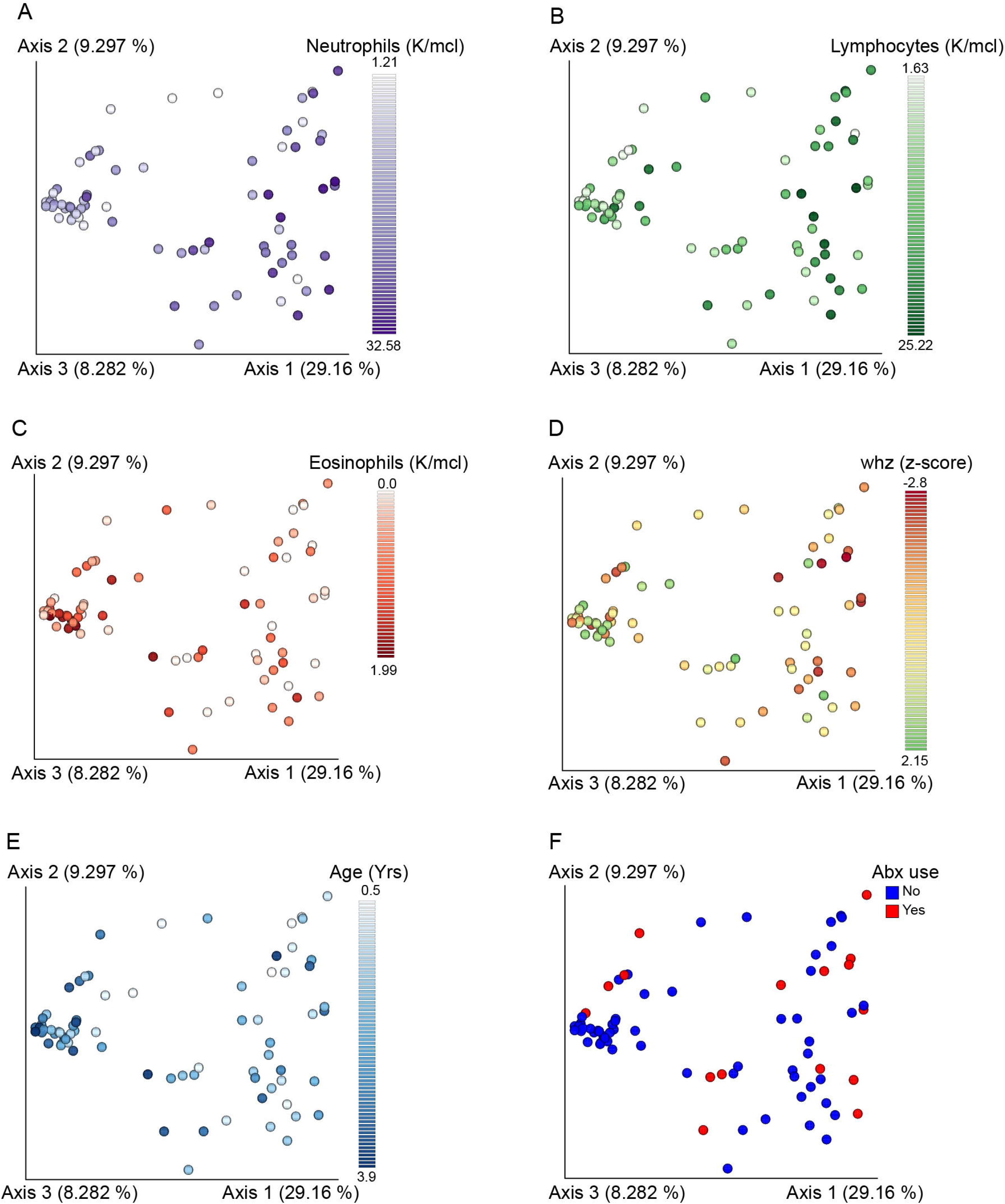
Beta diversity analysis of covariates that explain significant variation in the gut microbiota composition of children using Bray-Curtis distance. (**A-C**) PCoA plot shows clustering of Ugandan children by absolute neutrophils, lymphocytes, and eosinophils count, respectively. (**D-E**) PCoA plot shows clustering of Ugandan children by weight for height z-score (whz) and age, respectively. (**F**) PCoA plot shows clustering of Ugandan children by antibiotic use.

**Figure S7.**
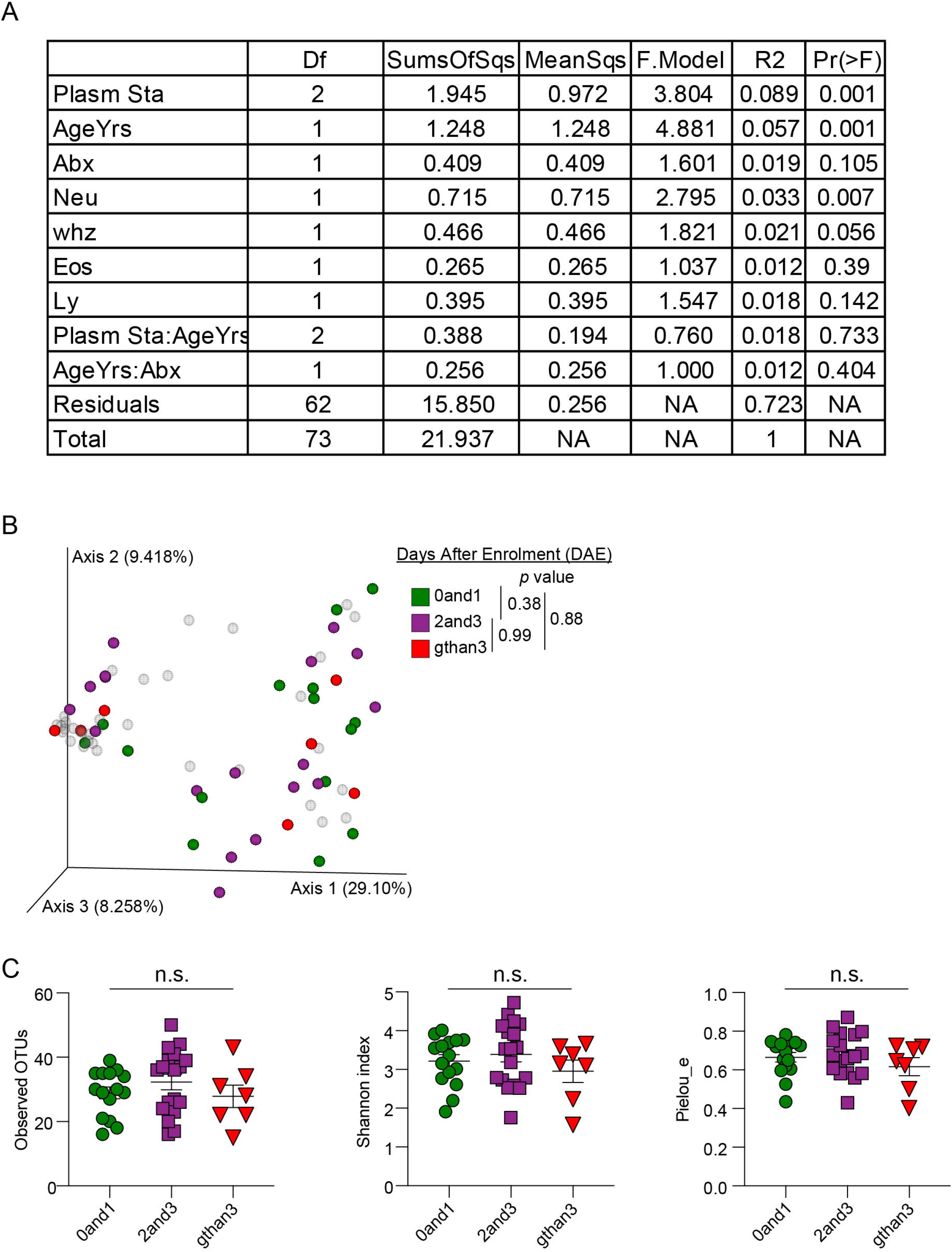
Stool bacteria beta diversity modeling output and diversity analysis of SMA children whose stool were collected at varying days post enrollment. (**A**) Beta diversity analysis were performed at 5000 sequencing depth. One individual from Pf Neg group was excluded because of low sequencing depth resulting in 27 samples. Modeling *Plasmodium* status (Pf Neg, Pf Pos and SMA) from Fig. 2F using “adonis” function of Vegan package accounting for age of infants, antibiotic use, absolute neutrophil and whz with formula, Beta diversity ~ PlasmSta * AgeYrs * Abx + Neutrophils + whz + Eosinophils + Lymphocytes. (**B** and **C**) Bacterial diversity analysis of only SMA children with varying stool collection days after the enrollment. (**B**) PCoA plot shows varying days of stool collection after the enrollment. Light shaded area are Pf Pos and Pf Neg samples. (**C**) Alpha diversity analysis. Data are means ± SEM. Statistical significance were analyzed by pairwise PERMANOVA (**B**) and pairwise Kruskal-Wallis test (**C**) (p < 0.05). n.s. = not significant.

**Figure S8.**
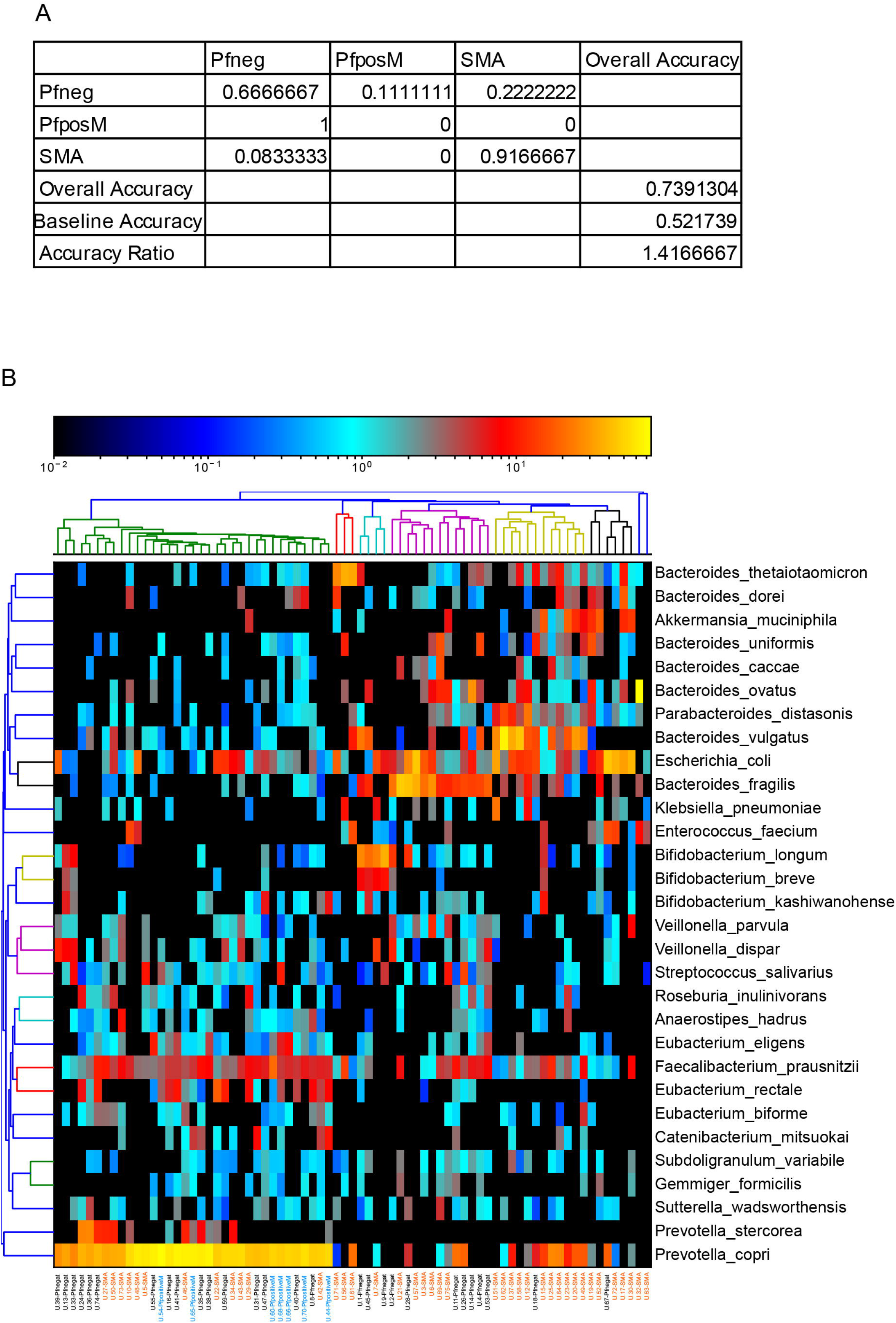
Accuracy of machine-learning model using random forest and taxonomic assignment. (**A**) Model accuracy of machine-learning using random forest. (**B**) Relative abundance of top 30 bacterial species. Hierarchical clustering of samples and features were performed using average linkage method.

**Figure S9.**
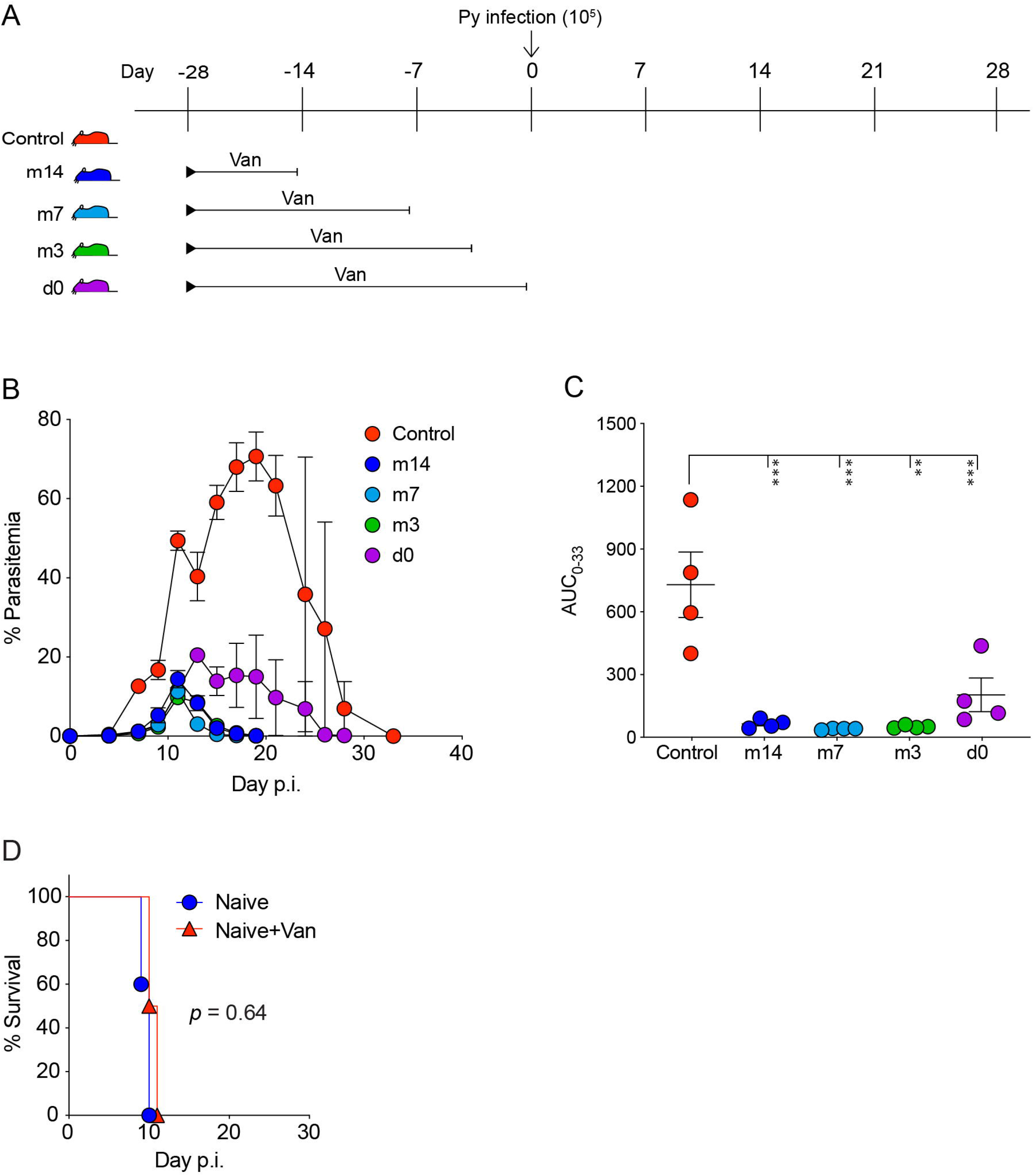
Vancomycin treatment provides protection against *P. yoelii* for weeks. (**A**) Mice were treated with vancomycin in drinking water starting 28 day before Py infection. Vancomycin treatment was stopped on the indicated days [-14 (m14), −7 (m7), −3 (m3), and 0 (d0)] before Py infection. Control mice received no vancomycin. N = 4 mice per group. (**B** and **C**) Parasitemia and AUC respectively. Data are means ± SEM. (**C**) One-way ANOVA with Tukey’s multiple comparisons test. Data are representative of multiple experiments. ** = *p* < 0.01, *** = *p* < 0.001. (**D**) Survival analysis of CR mice treated with from day 0 p.i. These mice were infected with *P. berghei* ANKA. N = 5 mice/group. Pairwise Log-rank (Mantel-Cox) test.

**Figure S10.**
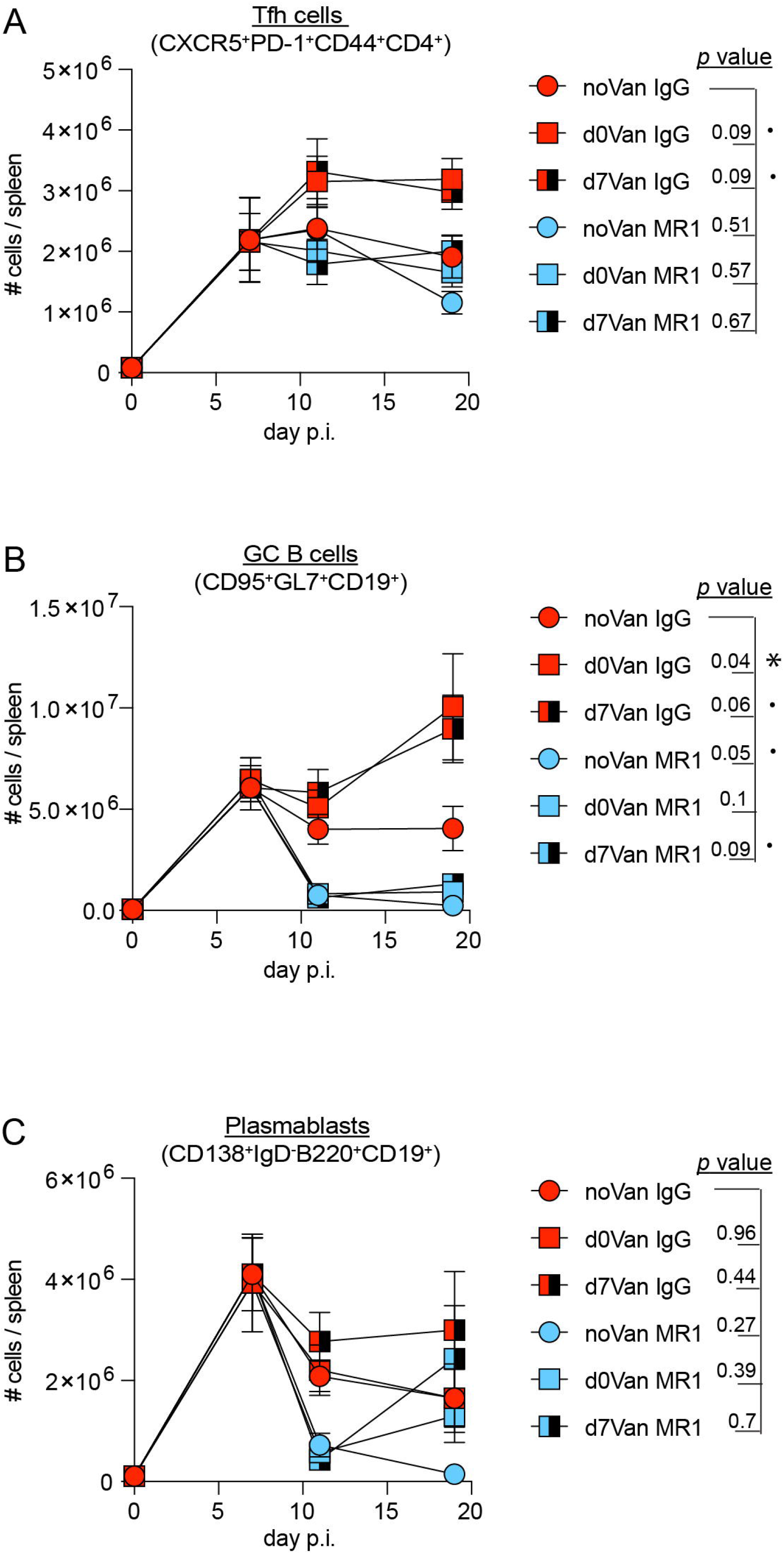
Cellular analysis following vancomycin and MR1 treatment. Cellular analysis from Fig. 6 is represented longitudinally. (**A**) Tfh cells (CXCR5^+^PD-1^+^CD44^+^CD4^+^) per spleen of mice harvested on day 0, 7, 11, and 19 p.i. (**B**) GC B cells (CD95^+^GL7^+^CD19^+^) per spleen of mice harvested on day 0, 7, 11, and 19 p.i. (**C**) Plasmablasts (CD138^+^lgD’B220^+^CD19^+^) per spleen of mice harvested on day 0, 7, 11, and 19 p.i. Data are means ± SEM. Statistical analyses were performed using two-way ANOVA adjusting for FDR corrected using two-stage step-up method of Benjamini, Krieger and Yekutieli. Only the comparisons with noVan IgG are shown. · = *p* < 0.1; * *p* < 0.05.

## Supplementary Table

**Sheet 1**: Metadata of Uganda participants.

